# Diffusion geometry and topology reveal Alzheimer’s proteomic states

**DOI:** 10.64898/2026.11.13.766457

**Authors:** Kelly Shahu, Michael W. Lutz, Abhay R. Moghekar, Richard J. O’Brien, Tananun Songdechakraiwut

**Author notes:** **Correspondence:** Michael W. Lutz, Departments of Neurology and Pathology, Duke University School of Medicine, Joseph and Kathleen Bryan Research Building, 311 Research Drive, Room 201-D, Durham, NC 27710, USA., Tananun Songdechakraiwut, Department of Computer Science, Duke University, Levine Science Research Center, 308 Research Drive, Room 112-D, Durham, NC 27708, USA. Data used in preparation of this article were obtained from the Alzheimer’s Disease Neuroimaging Initiative (ADNI) database (adni.loni.usc.edu). As such, the investigators within the ADNI contributed to the design and implementation of ADNI and/or provided data but did not participate in analysis or writing of this report. A complete listing of ADNI investigators can be found at: http://adni.loni.usc.edu/wp-content/uploads/how_to_apply/ADNI_Acknowledgement_List.pdf. “For the Alzheimer’s Disease Neuroimaging Initiative*”.

## Abstract

**INTRODUCTION:** Alzheimer’s disease (AD) exhibits substantial molecular heterogeneity. Identifying biologically distinct proteomic states may improve understanding of disease heterogeneity and support precision diagnostics.

**METHODS:** Cerebrospinal fluid proteomic data from ADNI were analyzed. Among 6,361 proteins tested between cognitively normal and AD groups, 407 remained significant after Benjamini–Hochberg correction (FDR < 0.05), with 404 retained after missingness filtering. Following preprocessing and outlier removal, 130 individuals were included. Diffusion geometry and persistent homology were applied to characterize intrinsic proteomic structure without predefined clusters. The framework was further evaluated in an independent mass spectrometry cohort.

**RESULTS:** Persistent homology revealed multiscale organization of the AD proteomic landscape. At the selected H₀ filtration scale, two dominant proteomic states (n = 71 and n = 42) contained most individuals, alongside smaller components. H₁ persistence identified nontrivial cyclic structure, indicating continuous higher-order organization beyond discrete state partitions. The dominant states exhibited distinct proteomic profiles and differed substantially in tau-related biomarkers. Elevated tau pathology in one state was reproduced in the independent cohort, whereas Aβ42 differences were less consistent across cohorts.

**DISCUSSION:** AD proteomic heterogeneity is characterized by multiscale organization combining dominant group structure with continuous higher-order relationships.

## 1 | BACKGROUND

Alzheimer’s disease (AD) is a complex and progressive neurodegenerative disorder and the leading cause of dementia in the elderly, posing a significant public health challenge [1]. AD is primarily characterized by memory loss and cognitive decline and is biologically defined by the deposition of amyloid beta (Aβ) and hyperphosphorylated tau tangles [2, 3, 4]. However, the pathogenesis of AD extends beyond Aβ and tau alone, and increasing evidence indicates substantial heterogeneity in disease presentation and progression [5, 6].

Evidence has demonstrated that Alzheimer’s disease exhibits heterogeneity across multiple domains, including clinical symptoms, neuroimaging findings, and molecular profiles [5, 6, 7]. In particular, increasing attention has focused on proteomic heterogeneity, as variation in protein expression patterns reflects underlying biological processes that may drive differences in disease mechanisms [8, 9]. In addition, the progression trajectories of clinical and pathological features differ across individuals [5]. However, the underlying biological mechanisms contributing to this heterogeneity remain poorly understood [6]. Elucidating proteomic heterogeneity may therefore provide critical insight into disease pathophysiology and facilitate the development of targeted therapeutic strategies.

Clinical subtypes of AD have been investigated based on variation in cognitive profiles, age of onset, and neuroimaging features [10]. Nevertheless, these classifications have primarily focused on macroscopic phenotypes and have shown variable findings across studies [6]. This variability may reflect the influence of multiple factors on clinical features and highlights the need to investigate biological subtypes defined at the molecular level.

Changes in cerebrospinal fluid (CSF) proteins directly reflect brain physiology and provide insight into molecular processes associated with disease progression [11, 12]. CSF proteomics captures a wide range of biological pathways, including neuroinflammation, synaptic function, and metabolic processes [13]. Previous studies using CSF proteomics have identified subgroups of individuals with distinct molecular signatures [11]. However, these studies have often relied on a limited number of proteins, which may restrict the ability to capture subtle molecular differences across subtypes.

In addition, neuropathological studies have identified distinct patterns of tau distribution, including typical AD, limbic-predominant AD, and hippocampal-sparing AD [14]. These patterns are associated with differences in clinical presentation and progression [15]. Genetic studies have also identified multiple risk loci, including APOE, which contributes significantly to disease risk [13]. However, genetic factors alone do not fully explain the heterogeneity observed in AD.

Recent studies have applied data-driven approaches integrating multimodal data to identify biologically defined subtypes [16, 6, 17]. Notably, prior work using large-scale CSF proteomics has identified multiple molecular subtypes of AD. Earlier studies reported up to five subtypes associated with distinct biological processes and genetic risk profiles [18]. More recent work using machine learning approaches, including non-negative matrix factorization and random forest models, identified three molecular subtypes characterized by differences in biological pathways and disease progression patterns [11]. These findings support the use of data-driven approaches to define molecular heterogeneity in AD, although the number and characteristics of subtypes vary across studies. These subtypes corresponded to relatively stable, intermediate, and rapidly progressing disease trajectories, highlighting the clinical relevance of molecular stratification. However, differences in clinical presentation and cognitive decline across these subtypes are often modest, limiting their clinical utility [6]. This limitation may result from the restricted resolution of previous approaches or the complexity of the underlying biological structure.

In this study, we applied a data-driven framework to identify molecular states of AD using CSF proteomics. To capture the intrinsic structure of the data, we used diffusion-based distances [19] and persistent homology [20, 21, 22]. This approach allows the identification of stable structures in high-dimensional data without specifying the number of AD subtypes a priori. We then examined whether the identified subtypes were associated with differences in biomarker profiles and clinical characteristics to better define their biological relevance.

## METHODS

### 2.1 | Data

Data used in the preparation of this article were obtained from the Alzheimer’s Disease Neuroimaging Initiative (ADNI) database (adni.loni.usc.edu). The ADNI was launched in 2003 as a public-private partnership, led by Principal Investigator Michael W. Weiner, MD. The primary goal of ADNI has been to test whether serial magnetic resonance imaging (MRI), positron emission tomography (PET), other biological markers, and clinical and neuropsychological assessment can be combined to measure the progression of mild cognitive impairment (MCI) and early Alzheimer’s disease (AD). For up-to-date information, see www.adni-info.org.

### 2.2 | Proteomic data preprocessing

Cerebrospinal fluid (CSF) proteomic data were obtained from the study cohort and processed using a standardized preprocessing pipeline to ensure data quality and comparability across individuals. For feature selection, proteins were considered for analysis only if at least eight non-missing observations were available in both cognitively normal (CN) and Alzheimer’s disease (AD) groups. To focus on biologically relevant variation, differential expression between CN and AD groups was assessed using non-parametric statistical methods [11]. Proteins were further filtered to retain those with ≤10% missing values in the clustering cohort, consistent with prior proteomic studies [1].

For each protein, differences between CN and AD groups were assessed using the Kruskal–Wallis test. Resulting p-values were corrected for multiple comparisons across all tested proteins using the Benjamini–Hochberg false discovery rate (FDR) procedure, and proteins with an FDR-adjusted p-value < 0.05 were retained for subsequent analyses.

Remaining missing values were imputed using median imputation, a conservative approach that preserves marginal distributions while minimizing the introduction of artificial variance. Protein expression values were subsequently standardized using z-score normalization to ensure that downstream analyses were not dominated by proteins with larger absolute abundance ranges and that all features contributed comparably to distance calculations. Prior to downstream modeling, outlier samples were identified and removed using an isolation forest approach [23], an unsupervised anomaly detection method based on recursive partitioning that has been shown to perform effectively in high-dimensional biological datasets, thereby reducing the influence of technical artifacts and extreme values on manifold construction [24].

### 2.3 | Replication cohort

An independent replication cohort was used to evaluate the robustness and generalizability of the identified proteomic state structure. Replication data were obtained from a cerebrospinal fluid (CSF) proteomic dataset collected at Johns Hopkins University, comprising 101 individuals: 40 cognitively normal (CN) individuals, 19 individuals with early mild cognitive impairment (EMCI), 21 with late mild cognitive impairment (LMCI), and 21 with pathological AD. Clinical diagnoses were assigned according to established criteria, including standardized cognitive assessments and biomarker-based classifications consistent with ADNI and NIA-AA recommendations.

CSF samples were collected via lumbar puncture under standardized conditions and processed using mass spectrometry–based proteomics, providing an orthogonal measurement platform relative to the aptamer-based SomaScan technology used in the discovery cohort. This distinction enables validation of proteomic state structure across fundamentally different proteomic acquisition modalities, thereby strengthening the biological interpretation and generalizability of the results.

Proteomic measurements were generated using data-independent acquisition (DIA) liquid chromatography–mass spectrometry (LC-MS/MS). Raw data were processed using Spectronaut (Biognosys), with peptide and protein identification controlled at a 1% false discovery rate. Protein quantification was performed using the MaxLFQ approach, and post-translational modification localization was incorporated where applicable. Quality control included removal of peptide outliers and filtering based on completeness across diagnostic groups.

Proteomic data underwent standardized preprocessing before the topological analysis. Protein intensities were log2-transformed to stabilize variance, and batch effects arising from plate, column, and FAIMS acquisition conditions were corrected using ComBat with non-parametric adjustments. Proteins with less than 25% completeness across diagnostic groups were excluded, and remaining missing values were imputed using a random forest-based approach (missForest).

To ensure comparability with the discovery analysis, the cognitively normal group was used as the reference population for normalization and disease-associated feature selection. Protein feature selection was performed by comparing the 40 cognitively normal individuals with the 21 pathological AD individuals using the Kruskal–Wallis test, followed by Benjamini–Hochberg correction for multiple testing. This procedure identified 93 proteins at FDR < 0.05, which were retained for the subsequent subtype analysis.

Subtype identification was performed separately within the disease-spectrum cohort, comprising individuals with EMCI (n = 19), LMCI (n = 21), and pathological AD (n = 21), for a total of 61 individuals. Cognitively normal individuals were therefore used as a reference population for normalization and feature selection but were not included in the disease-subtyping analysis. Within the disease-spectrum cohort, proteins with more than 10% missing values were excluded, individuals with less than 80% protein completeness were removed, and remaining missing values were imputed using the protein median. Protein values were subsequently standardized across the disease-spectrum cohort. Isolation Forest with a contamination parameter of 0.05 was then applied as an outlier quality-control step, removing three individuals and leaving 58 individuals for diffusion-distance construction and persistent-homology analysis.

### 2.4 | Construction of diffusion-based distance

To capture the intrinsic geometry of the high-dimensional proteomic space, pairwise relationships between individuals were modeled using a diffusion-based distance rather than conventional Euclidean distance [25].

Euclidean distance assumes that data lie in a linear space and treats all dimensions as independent and equally informative. However, proteomic data are inherently nonlinear and structured by complex biological processes, including co-regulation, pathway interactions, and latent disease trajectories. In these settings, Euclidean distance is sensitive to noise and can obscure meaningful relationships by overemphasizing high-dimensional variance.

To address these limitations, a diffusion geometry framework was employed [25, 19]. Specifically, a graph was constructed in which nodes represent individuals and edges encode local similarities between them, defined by whether two individuals are mutual k-nearest neighbors in the protein expression space.

Diffusion distances were constructed from a k-nearest-neighbor graph (k = 15) based on cosine distances between proteomic profiles. Edge weights were defined using a Gaussian kernel, 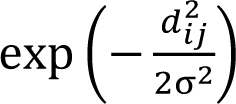, where the bandwidth (σ) was set adaptively to the median distance across all k-nearest-neighbor edges. The symmetric affinity matrix was row-normalized to obtain a Markov transition matrix (P), without additional α-density normalization. A diffusion time of (t=3) was used, and diffusion distances between individuals were calculated as the Euclidean distances between the corresponding rows of (P^3^). This approach captures the underlying manifold structure of the data, is robust to noise, and emphasizes global organization while preserving local neighborhoods.

By integrating information over multiple steps of the diffusion process, this metric effectively denoises the data and enhances biologically meaningful structure, enabling more accurate identification of patient subgroups compared to Euclidean-based approaches [19].

### 2.5 | Persistent homology analysis

The diffusion-based distance matrix constructed in Section 2.4 serves as the input for subsequent topological analysis. By encoding multi-step relationships between individuals, this representation provides a geometry-aware metric that captures both local and global structure of the data. Persistent homology is then applied to this distance matrix to identify stable topological features that reflect intrinsic subgroup organization.

Topological data analysis (TDA) [20, 21] was used to characterize the shape of the proteomic data and to identify intrinsic subgroup structure without imposing parametric assumptions. Persistent homology was computed on the diffusion-based distance matrix using a Vietoris–Rips filtration.

In this framework, a filtration parameter ε is gradually increased, and simplicial complexes are constructed by connecting points whose pairwise distance is below ε. Topological features, including connected components (H₀), loops (H₁), and higher-dimensional voids (H₂), are tracked across the filtration. Each feature is associated with a birth and death scale, and its persistence (lifetime) reflects its stability across scales.

Persistent features, defined by long lifetimes, are interpreted as robust structural characteristics of the data, whereas short-lived features are typically attributable to noise [26]. This multiscale representation provides a principled way to distinguish meaningful biological structure from stochastic variation, a key advantage over traditional clustering methods that operate at a single resolution [25].

### 2.6 | Subtype identification using H₀ persistence

Subtype identification was performed using zero-dimensional persistent homology (H₀), which tracks the formation and merging of connected components as the filtration parameter increases [27]. For a Vietoris–Rips filtration constructed from a pairwise distance matrix, this H₀ connectivity hierarchy is mathematically equivalent to the hierarchy produced by single-linkage clustering on the same distances [28]. Accordingly, we use H₀ persistence here as a topological representation of the connected-component hierarchy rather than as a clustering procedure fundamentally distinct from single-linkage hierarchical clustering.

Death times from the H₀ persistence diagram were extracted and sorted to characterize the scales at which connected components merged. Candidate thresholds were defined at midpoints between consecutive death times, and connected components were computed at each threshold. Candidate solutions were retained when at least two components contained a minimum of 10 individuals. This minimum-size criterion was introduced to restrict downstream subtype characterization to sufficiently represented groups and to avoid interpreting very small components as stable patient subtypes.

Among the retained candidate solutions, the filtration threshold was selected by maximizing the silhouette score calculated from the diffusion-distance matrix using the corresponding connected-component assignments. We therefore emphasize that the final threshold, and consequently the reported number of candidate subtypes, was selected using a non-topological internal clustering criterion rather than determined by persistent homology alone. Diagnostic labels and clinical outcomes were not used during threshold selection; thus, although the selection criterion is non-topological, it remains unsupervised with respect to clinical information.

Because H₀ persistence and single-linkage clustering encode the same connected-component hierarchy, the role of the H₀ analysis in this study is specifically to characterize the hierarchical organization of the diffusion-derived patient space and provide a consistent framework in which this connected-component structure can subsequently be considered alongside higher-dimensional topological features, particularly H₁ cycles. More generally, persistence diagrams provide a stable representation of topological structure under perturbations of the underlying data [29].

### 2.7 | Stability assessment via bootstrap resampling

To evaluate the robustness of the identified subtypes, a bootstrap resampling procedure was performed. The persistent homology–based clustering pipeline was applied to multiple resampled datasets generated by sampling individuals with replacement.

For each bootstrap iteration, subtype assignments were compared to those obtained from the full dataset using the adjusted Rand index (ARI), which quantifies agreement between clustering solutions while accounting for chance. The distribution of ARI values across bootstrap iterations was used to assess clustering stability.

High ARI values indicate that the identified subtype structure is reproducible under perturbations of the data, supporting its robustness and reducing the likelihood that results are driven by sampling variability.

### 2.8 | Null model testing of topological signal

To determine whether the observed topological structure exceeded that expected from the multivariate dependence structure of the proteomic data, we constructed a Gaussian-copula null model. Unlike independent feature permutation, which disrupts cross-protein dependencies, the Gaussian-copula framework preserves the empirical marginal distributions of individual proteins while approximating the dependence structure among proteins.

Protein measurements were transformed to Gaussian rank coordinates, and the corresponding latent correlation matrix was estimated from the observed data. Synthetic datasets were then generated from the fitted multivariate Gaussian copula and mapped back to the empirical marginal distribution of each protein. For each null realization, the complete diffusion-distance construction and persistent-homology analysis were repeated using the same parameters as for the observed data.

A total of 1,000 Gaussian-copula null realizations were generated. For H₀, we evaluated the largest and second-largest finite persistence values and the persistence gap between the second- and third-ranked H₀ features. Because finite H₀ death times correspond to minimum-spanning-tree edge weights, these statistics quantify prominent scales in the connected-component hierarchy.

For H₁, we evaluated the largest and second-largest persistence values, the persistence gap between the second- and third-ranked features, and total H₁ persistence. Empirical one-sided p-values were calculated by comparing each observed statistic with its corresponding null distribution using a finite-simulation correction. This framework tests whether specific H₀ or H₁ features are more pronounced than expected from correlated multivariate proteomic variation alone.

### 2.9 | Subtype refinement and downstream analyses

To facilitate downstream analyses and improve interpretability, subtype assignments were refined following data-driven identification of clusters using H₀ persistence. Candidate solutions were retained when at least two components exceeded the predefined minimum size threshold, and the operational threshold was selected using silhouette-based evaluation. From this solution, individuals belonging to the two largest components were retained for downstream characterization (n = 113), while 17 individuals in components containing fewer than 10 individuals were excluded from downstream group-level analyses because of limited sample size.

Associations between subtype membership and genetic or clinical variables were assessed using both unadjusted and adjusted models for age and sex. Categorical associations were evaluated using chi-square or Fisher’s exact tests as appropriate, while logistic regression models were used to estimate adjusted effects, controlling for covariates such as age and sex.

Genetic risk was operationalized using APOE ε4 carrier status, derived from available genotype data. Model coefficients were exponentiated to obtain odds ratios, facilitating interpretation of effect sizes.

### 2.10 | Pathway enrichment analysis

To delineate the biological processes underlying subtype-specific proteomic differences, we first performed differential protein expression analysis between the two dominant proteomic subtypes. Statistical significance was assessed using a two-sided Welch’s t-test (unequal variance), with a threshold of p < 0.01.

Proteins identified as significantly differentially expressed were subsequently used as input for pathway enrichment analysis using the g:Profiler framework [30]. Enrichment significance was evaluated using overrepresentation analysis with multiple testing correction, comparing the observed protein set to annotated biological pathways.

All analyses were performed with the organism parameter set to Homo sapiens (“hsapiens”), ensuring that functional annotations were derived from human-specific databases, including Gene Ontology and Reactome [12, 31].

### 2.11 | Pathway similarity network

To further characterize relationships among enriched biological processes, we constructed a pathway similarity network. In this representation, each node corresponds to an enriched pathway, annotated by its enrichment significance and gene ratio.

Pairwise similarity between pathways was defined using a combination of semantic and quantitative criteria. Pathway names were decomposed into informative keywords after removing non-informative terms (e.g., “process,” “regulation”), and pathways sharing one or more keywords were considered functionally related. Additional weighting was applied for pathways originating from the same annotation source or exhibiting similar gene ratios.

Edges were retained when the combined similarity score exceeded a predefined threshold, ensuring that the network emphasized robust and biologically meaningful relationships. The resulting network was visualized using a force-directed layout, in which spatial proximity reflects pathway similarity. Node size was scaled according to gene ratio, and node color encoded enrichment significance (−log10 *p*-value).

## RESULTS

### 3.1 | Participant characteristics and proteomic feature selection

The study cohort consisted of individuals from the ADNI dataset with available baseline cerebrospinal fluid (CSF) proteomic measurements. A total of 708 participants and 6,361 proteins were initially included.

To identify proteins associated with disease status, non-parametric testing was performed comparing cognitively normal (CN) and Alzheimer’s disease (AD) groups.

Of the 6,361 proteins tested, 559 met the nominal significance threshold (p < 0.01); following Benjamini–Hochberg correction for multiple testing, 407 proteins remained significant at FDR < 0.05 and were retained for downstream analyses.

Clustering analyses were subsequently restricted to individuals diagnosed with AD or mild cognitive impairment (MCI), yielding 137 participants with complete proteomic measurements across the selected features. After applying missingness filtering and quality control procedures, 404 proteins remained.

Outlier detection using an isolation forest algorithm identified 7 individuals as anomalous, which were removed from further analysis. The final analytical cohort consisted of 130 participants with high-quality proteomic profiles.

### 3.2 | Identification and topological characterization of proteomic states

To characterize the intrinsic structure of the proteomic data, diffusion-based distances were computed and used to construct a Vietoris–Rips filtration. Persistent homology of the resulting distance matrix revealed 129 H₀ death events, corresponding to the progressive merging of connected components across filtration scales. The associated minimum spanning tree (MST), whose edge weights correspond to finite H₀ death times, was used to characterize this hierarchical connectivity [13].

Examination of the MST edge-weight distribution identified a largest gap whose midpoint occurred at ε = 0.0416 [25]. At this filtration scale, two connected components remain, corresponding to a stable region of the Betti-0 curve (Figure 1A). This scale was used only to characterize the dominant large-scale organization of the proteomic space and was not used to assign the proteomic states analyzed downstream.

**FIGURE 1.**
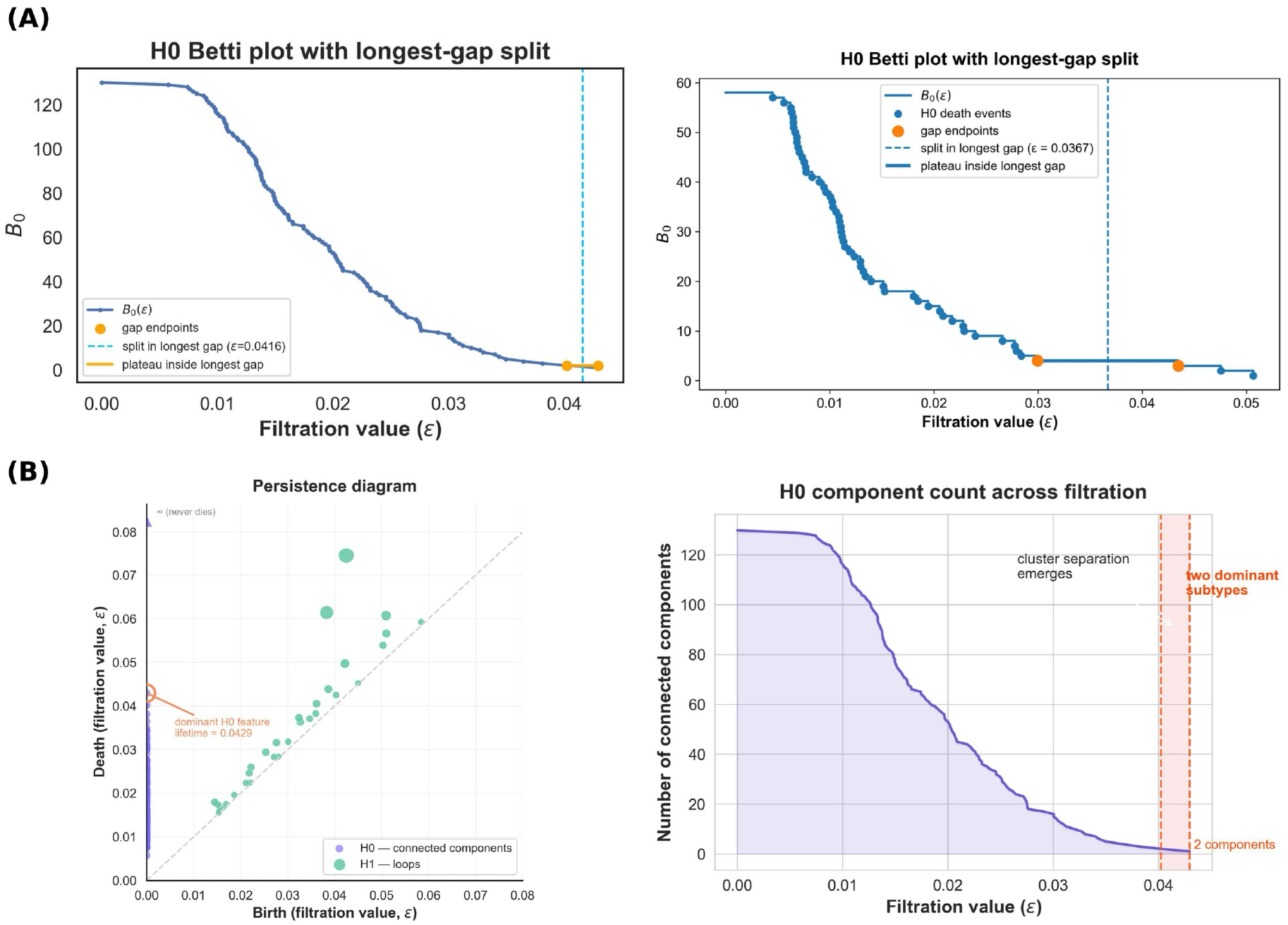
Topological evidence of dominant proteomic subtype structure in Alzheimer’s disease cohorts. **(A)** Betti-0 analysis of proteomic structure in the discovery and replication cohorts. Betti-0 curves show the number of connected components (H₀) as a function of the filtration parameter (*ε*) for the discovery (left) and replication (right) cohorts. Points denote H₀ death events corresponding to merge operations in the minimum spanning tree (MST). The largest gap between consecutive MST edge weights is highlighted, and the vertical dashed line indicates the midpoint of this gap, illustrating a prominent coarse-scale transition in the connected-component hierarchy. This threshold is shown descriptively and was not used for downstream state assignment. Across both cohorts, the H₀ structure demonstrates multiscale changes in connectivity as the filtration parameter increases. **(B)** Persistence-based visualization of subtype separation in the discovery cohort. The persistence diagram (left) summarizes both H₀ and H₁ topological features. H₀ features represent connected components, with the dominant H₀ feature corresponding to the longest-lived connected component split that separates the major proteomic subtypes. H₁ features represent persistent loop-like structures (cycles) in the patient similarity network, capturing higher-order organization beyond connected components. The H₀ filtration curve (right) illustrates the evolution of connected components across the filtration process and highlights the emergence of two dominant connected components, supporting robust topological separation between the major proteomic subtypes.

For state assignment, candidate thresholds identified from the H₀ lifetime spectrum were evaluated separately. Among these thresholds, ε = 0.0335 maximized the silhouette score under the predefined minimum cluster-size criterion. At ε = 0.0335, the data partitioned into multiple connected components, with two dominant clusters containing the majority of individuals (n = 71 and n = 42), alongside several smaller components containing fewer than 10 patients. Subsequent biological and clinical analyses therefore focused on these two largest states, comprising 113 individuals.

Importantly, the two-component regime observed at the coarser ε = 0.0416 scale should not be interpreted as evidence that the proteomic landscape is fundamentally bipartite. H₀ describes a multiscale connectivity hierarchy in which multiple components present at finer filtration values progressively merge as ε increases. The two-component regime therefore represents one prominent level of this hierarchy, while the ε = 0.0335 threshold provides the operational subtype definition used for downstream analyses. This interpretation is further complemented by the H₁ analysis below, which captures continuous and nonlinear relationships that cannot be represented by connected-component partitions alone.

### 3.3 | Cycle analysis of proteomic structure

To further characterize higher-order organization beyond connected components, we examined one-dimensional homological features (H₁), which capture loop-like structure in the underlying similarity graph. In contrast to H₀ components, which reflect disconnected regions of the space, H₁ features arise when connectivity forms nontrivial cycles that cannot be reduced to a tree-like structure. These cycles were identified from the persistence diagram (Figure 1B) computed on the patient–patient distance matrix, with each H₁ class corresponding to a topological loop that persists across a range of filtration values.

We observed two leading H₁ features with persistence values of 0.0321 and 0.0231, respectively. To determine whether these features represented structure beyond that expected under a dependence-preserving null model, we compared the observed H₁ persistence spectrum with 1,000 Gaussian-copula null realizations. The largest H₁ persistence did not reach statistical significance relative to the null distribution (observed = 0.0321; null mean = 0.0210; empirical p=0.085). In contrast, the second-largest H₁ feature showed significantly greater persistence than expected under the Gaussian-copula null (observed = 0.0231; null mean = 0.0133; empirical p=0.026). We additionally evaluated the persistence gap between the second- and third-ranked features. The observed gap was 0.0133, approximately fourfold greater than the null mean of 0.00331, and exceeded 98.4% of null realizations (empirical p=0.017). This significant separation supports the presence of two leading persistent cycles that are distinguishable from the remaining H₁ persistence spectrum. The most persistent H₁ feature had birth ε≈0.0425, death ε≈0.0745, and persistence ≈0.0321, with a representative cocycle involving 36 patients. The second-ranked H₁ feature had birth ε≈0.0383, death ε≈0.0614, and persistence ≈0.0231, with a representative cocycle involving 11 patients. To explore whether the representative cycles corresponded to clinically distinct patient profiles, we compared patients associated exclusively with each of the two leading cocycle representatives, excluding the single patient shared between them. This yielded 35 patients associated exclusively with the first representative and 10 with the second. After false-discovery-rate correction across the evaluated clinical variables, the groups differed significantly in APOE ε4 dosage (FDR = 0.012) and CSF amyloid-β levels (FDR = 0.012), whereas differences in tau did not remain significant after multiple-testing correction (FDR = 0.052). APOE ε4 carrier prevalence was also higher among patients associated exclusively with the second representative (100% versus 62.9%; Fisher’s exact p=0.042). Given the small sample size of the second group and the non-uniqueness of cocycle representatives, these clinical associations are interpreted as exploratory rather than as evidence of discrete patient subtypes. These cycles are visualized in Figure 2, where nodes represent patients and edges encode pairwise similarity below the filtration threshold. The highlighted red edges correspond to representative cocycles selected from an equivalence class of cycles defining the same topological feature, providing an interpretable visualization of the underlying loop geometry. Importantly, these cocycles are not unique; rather, they represent one possible realization of a broader class of equivalent cycles, and should therefore be interpreted as evidence of underlying topological loops rather than literal geometric structures.

**FIGURE 2.**
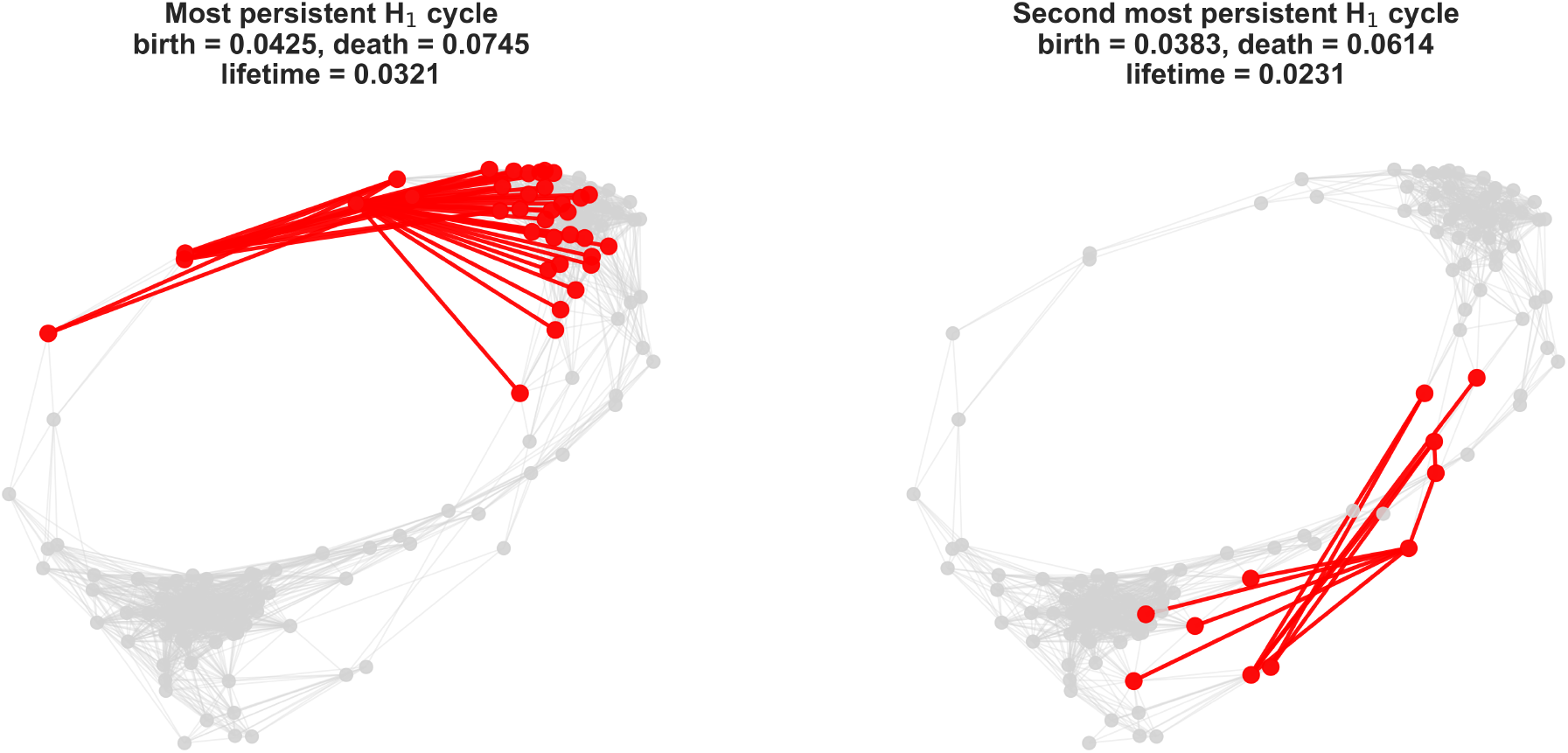
Cycle structure of the proteomic similarity graph revealed by persistent homology. Left: Spring layout visualization of the most persistent H₁ feature, showing a global loop spanning a subset of patients within a densely connected region of the graph. Nodes represent individuals, and edges indicate pairwise similarity below the filtration threshold. The highlighted red edges correspond to a representative cocycle, illustrating the presence of nontrivial cyclic connectivity in the patient similarity structure. Right: Visualization of the second most persistent H₁ feature, corresponding to a more localized loop involving a smaller subset of patients. This structure reflects a cycle within a restricted region of the proteomic space. Together, these H₁ features indicate that, beyond the dominant hierarchical subtype organization, the proteomic landscape exhibits higher-order topology with loop-like structure, suggesting the presence of continuous relationships among subsets of patients.

Visualization of representative cocycles on the filtration graph confirms that these features reflect nontrivial cyclic connectivity patterns rather than isolated clusters. Importantly, these H₁ features arise within the broader hierarchical organization identified by the H₀ analysis, in which the cohort separates into two dominant components. The presence of persistent cycles suggests that, in addition to this hierarchical organization, the proteomic landscape contains continuous relationships among patients that are not fully captured by discrete partitions alone. Biologically, these loop-like structures indicate that patient similarity is not strictly partitioned into isolated subtypes, but instead includes nonlinear transitions and overlapping relationships that may reflect gradual changes in molecular states or shared pathological processes. Visualization of representative cocycles on the filtration graph illustrates that the leading H₁ features reflect cyclic connectivity patterns that are not captured by connected-component structure alone. The Gaussian-copula analysis provides statistical support for the second-ranked H₁ feature and for the separation of the two leading cycles from the remainder of the persistence spectrum. These findings suggest that the proteomic landscape contains higher-order organization beyond the connected-component hierarchy captured by H₀, although the biological interpretation of individual cocycle representatives should remain cautious because representatives of a given homology class are not unique.

### 3.4 | Geometric evidence of multiscale proteomic organization

To further examine the multiscale structure identified by persistent homology, we visualized the organization of the proteomic space across the filtration. The connectivity pattern changed progressively with filtration scale, with multiple components at finer scales merging into a smaller number of larger structures as ε increased. Within this hierarchy, a regime containing two dominant connected components was evident at a coarser scale.

Importantly, this two-component regime represents one scale of the connectivity hierarchy rather than a strict bipartition of the full proteomic landscape. At the downstream analysis threshold, additional smaller components remained, while H₁ analysis identified cyclic relationships extending beyond discrete connected-component assignments. A comparable large-scale connectivity pattern was observed in the independent replication cohort, suggesting that aspects of this multiscale organization are reproducible across cohorts and proteomic platforms. Figure 3 visualizes this scale-dependent organization, including dominant groups, smaller components, and finer internal structure.

**FIGURE 3.**
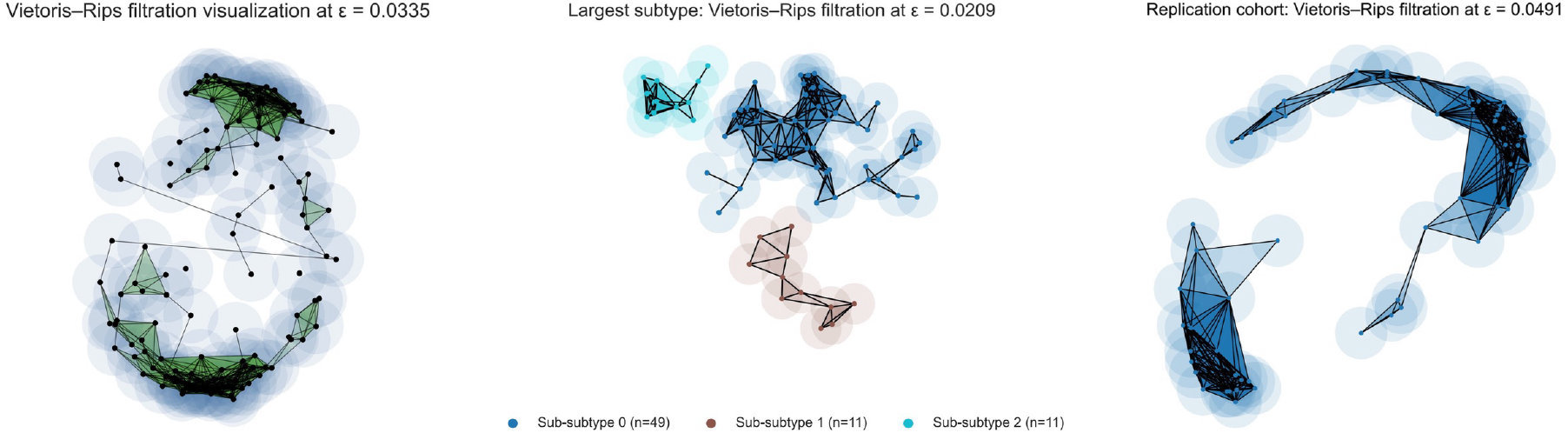
Multi-scale Vietoris–Rips filtration visualizations of proteomic similarity structure across discovery and replication cohorts. Left: Vietoris–Rips filtration of the full discovery cohort at the silhouette-selected operational threshold (ε = 0.0335), illustrating the emergence of two dominant connected components (H₀), consistent with the primary subtype separation. Center: Filtration restricted to the largest subtype, revealing three sub-subtypes at a finer scale (ε = 0.0209), corresponding to internal H₀ splits within the dominant cluster. Right: Replication cohort filtration at its silhouette-selected operational threshold (ε = 0.0491), demonstrating a consistent large-scale organization with a dominant connected structure and smaller peripheral components. In all panels, nodes represent patients, edges indicate pairwise similarity below ε, and filled triangles denote higher-order simplices (2-simplices), highlighting local geometric connectivity. The persistence of these structures across scales and cohorts supports a hierarchical organization of Alzheimer’s disease heterogeneity.

To evaluate whether the observed H₀ organization exceeded that expected from the multivariate dependence structure of the proteomic data, we compared the observed H₀ persistence statistics with 1,000 Gaussian-copula null realizations. None of the evaluated H₀ statistics differed significantly from their corresponding null distributions. The largest H₀ persistence was 0.0506 compared with a null mean of 0.0457 (empirical p = 0.170), while the second-largest H₀ persistence was 0.0423 compared with a null mean of 0.0419 (empirical p = 0.433). Similarly, the persistence gap between the second- and third-ranked H₀ features was 0.00226 compared with a null mean of 0.00249 (empirical p = 0.399).

These results indicate that the prominent scales observed in the H₀ connected-component hierarchy are not more pronounced than expected under a null model that preserves the marginal protein distributions and approximates their multivariate dependence structure. Accordingly, the H₀-defined groups are interpreted as a useful scale-dependent partition of the proteomic landscape rather than as statistical evidence, by themselves, for intrinsically discrete patient subtypes. Their biological relevance is therefore evaluated through downstream molecular and clinical characterization, replication, and stability analyses rather than inferred from H₀ significance alone.

To further assess robustness, we performed 1,000 bootstrap resampling iterations and compared proteomic state assignments with the corresponding original assignments using the adjusted Rand index (ARI). Restricting this analysis to the two dominant connected components (n = 113), proteomic state assignments showed high reproducibility across bootstrap samples (mean ARI = 0.858, SD = 0.158; 2.5th–97.5th percentile interval: 0.476–1.000), demonstrating that state assignments at the selected filtration scale were reproducible under resampling.

### 3.5 | Functional enrichment analysis of proteomic state-associated proteins

To delineate the biological processes underlying proteomic state differences, we performed differential protein expression analysis between the two dominant proteomic states, followed by pathway enrichment analysis. Statistical significance for differential expression was assessed using a two-sided Welch’s t-test (p < 0.01), and pathway enrichment was evaluated using overrepresentation analysis with multiple testing correction [30, 12].

At the individual protein level, several of the top differentially expressed proteins are functionally linked to synaptic and neuronal processes. Notably, proteins such as DLG2 and LRTM2, which are associated with synaptic organization and cell junction assembly, and EPHA5, a receptor involved in axon guidance and synaptic connectivity, were among the most significantly altered. Additional proteins, including STMN1 and STMN2, which regulate microtubule dynamics and neuronal structure, further support the presence of coordinated alterations in neuronal signaling pathways [12]. These observations indicate that proteomic state differences are reflected not only at the pathway level but also in specific proteins with established roles in synaptic biology.

Consistent with these protein-level findings, enrichment analysis revealed that proteins distinguishing Subtype 1 from Subtype 0 were significantly overrepresented in pathways related to synaptic structure, neuronal signaling, and cell–cell communication (Figure 4A) [12, 9]. The most significant terms included synapse, cell junction, synaptic signaling, and trans-synaptic communication. Importantly, enrichment was not limited to isolated pathways but was consistently observed across multiple related functional categories, indicating a coordinated perturbation of neuronal communication systems. The convergence of enrichment signals across functionally overlapping pathways suggests that differences between the two proteomic states reflect broad disruption of synaptic processes rather than single-pathway effects [4, 10]. The complete list of enriched pathways is provided in Table S1.

**FIGURE 4.**
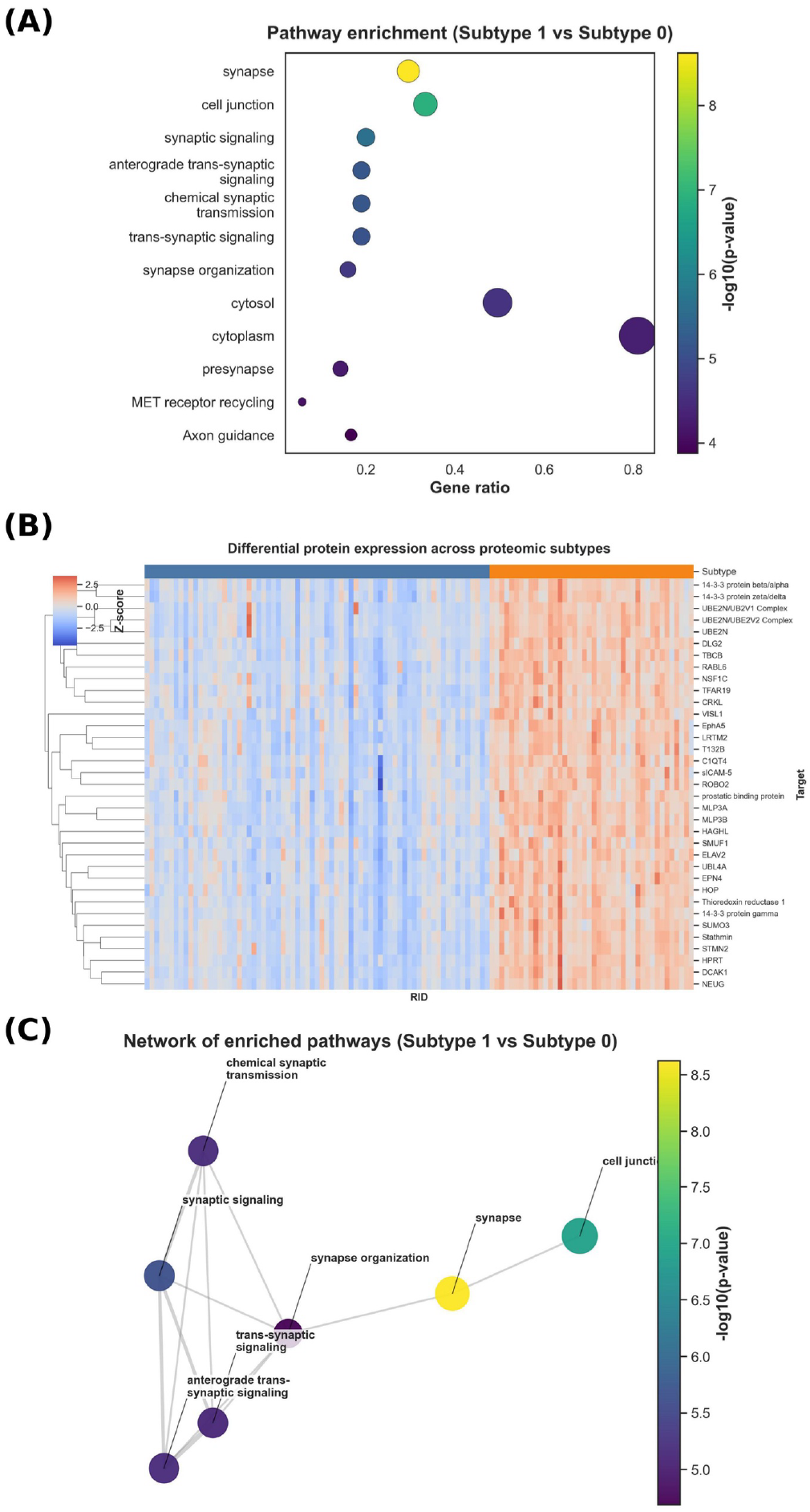
Biological pathway organization associated with dominant proteomic subtypes in Alzheimer’s disease. **(A)**Pathway enrichment analysis distinguishing the two dominant proteomic subtypes in the discovery cohort. Differentially expressed proteins between subtypes were enriched for pathways related to synaptic structure, neuronal signaling, cell junction organization, and trans-synaptic communication. Points are sized according to gene ratio (proportion of subtype-associated proteins overlapping each pathway) and colored by statistical significance (−log10 p-value), indicating coordinated alterations in synaptic and neuronal processes between subtypes. **(B)** Clustered heatmap of the top differentially expressed proteins between Subtype 1 and Subtype 0 in the discovery cohort. Rows are proteins (hierarchically clustered) and columns are individuals, shown as z-scored expression; the colored bar above the heatmap indicates subtype membership. Proteins show coordinated expression patterns within each subtype, with higher expression in Subtype 1. **(C)** Network organization of enriched biological pathways. Nodes represent enriched pathways, with node size proportional to the number of overlapping proteins and node color indicating enrichment significance (−log10 p-value). Edges reflect similarity between pathways based on shared functional annotations. The resulting network reveals a densely interconnected module centered on synaptic signaling and neuronal communication pathways, suggesting convergence of subtype-associated proteins onto coherent biological programs.

At the protein level, a clustered heatmap of the top differentially expressed proteins demonstrated clear separation between Subtype 0 and Subtype 1 across individuals (Figure 4B). Proteins exhibited coordinated expression patterns within each state, forming distinct profiles. This structured organization supports the presence of state-associated proteomic signatures and indicates that the observed differences involve coordinated variation across multiple proteins rather than isolated protein changes [8].

To further characterize the organization of these biological signals, we examined a network of enriched pathways based on functional similarity (Figure 4C). Enriched pathways formed a densely interconnected network centered on synaptic and neuronal processes. Highly significant pathways, including synapse and cell junction, occupied central positions within the network, while related signaling pathways formed tightly connected clusters. This network structure indicates that proteins differentiating Subtype 0 and Subtype 1 converge onto coherent and biologically related functional modules, rather than representing independent enrichment signals [30, 9].

Taken together, these findings indicate that Subtype 0 and Subtype 1 represent proteomic states characterized by distinct molecular programs, with prominent differences involving synaptic and neuronal processes. The consistency of these patterns across individual proteins, pathway enrichment, and network structure supports coordinated differences in neuronal signaling-related proteomic programs between the two states [18].

### 3.6 | Subtype characterization

Differential protein expression analysis revealed a marked asymmetry between the two proteomic states. CSF biomarker comparisons showed that Subtype 1 exhibited significantly higher levels of total tau (p = 6.84 × 10⁻¹⁴) and phosphorylated tau (p = 5.81 × 10⁻¹⁴), along with a smaller but statistically significant difference in Aβ42 (p = 3.00 × 10⁻²) (Figure S1A).

APOE ε4 carrier frequency differed significantly across controls and the two proteomic subtypes (χ² = 57.05, df = 2, p = 4.10 × 10⁻¹³). APOE ε4 carriage was observed in 24.6% of controls, 63.4% of Subtype 0, and 78.6% of Subtype 1. Relative to controls, both Subtype 0 (OR = 5.32, Fisher’s exact p = 2.43 × 10⁻⁸) and Subtype 1 (OR = 11.27, Fisher’s exact p = 1.63 × 10⁻¹⁰) showed significantly greater odds of APOE ε4 carriage. These associations remained significant after adjustment for age and sex (Subtype 0 vs controls: adjusted OR = 5.64, 95% CI 3.06–10.39, p = 2.77 × 10⁻⁸; Subtype 1 vs controls: adjusted OR = 11.36, 95% CI 4.95–26.07, p = 1.00 × 10⁻⁸). Although APOE ε4 carriage was numerically more frequent in Subtype 1 than in Subtype 0 (78.6% vs 63.4%), the direct subtype comparison did not reach statistical significance (OR = 2.12, Fisher’s exact p = 0.098). This difference remained non-significant after adjustment for age and sex (adjusted OR = 2.00, 95% CI 0.80–4.99, p = 0.136).

At the proteomic level, the volcano plot demonstrated that many of the most significant proteins showed both large effect sizes and strong statistical support, indicating coordinated shifts rather than isolated changes (Figure S1B). Notably, proteins such as DLG2, NRGN (assay identifier NEUG), DCLK1 (assay identifier DCAK1), and MAP1LC3A (assay identifier MLP3A) were among the most strongly upregulated in Subtype 1, whereas a smaller subset, including COQ7 and IFNL4, showed higher expression in Subtype 0. Consistent with these findings, the co-expression network in Subtype 1 revealed a densely connected core of proteins with high correlation structure, suggesting that subtype-associated differences are driven by coordinated biological programs rather than independent protein-level variation (Figure S1C). Together, these results indicate that Subtype 1 is characterized by a distinct molecular profile marked by widespread proteomic upregulation and increased tau pathology, supporting the presence of biologically differentiated disease states across subtypes. The complete differential protein expression results are provided in Table S2.

### 3.7 | Replication cohort

To assess the robustness of the identified subtype structure, the full analytical pipeline was applied to an independent replication cohort comprising 101 patients and 3,202 proteins. While the same preprocessing, feature selection, and topological analysis framework was used, the replication cohort was obtained using a different proteomics platform (mass spectrometry) compared to the SOMAscan-based discovery cohort, providing an additional level of validation across assay technologies.

Consistent with the discovery cohort, persistent homology of the diffusion-based distance matrix identified two dominant connected components. Restricting the analysis to these components yielded 53 individuals, partitioned into Subtype 0 (n = 34) and Subtype 1 (n = 19). Bootstrap analysis across 1,000 resampling iterations further supported the reproducibility of this partition (mean ARI = 0.762, SD = 0.248; median ARI = 0.849; 2.5th–97.5th percentile interval: 0.208– 1.000). Although variability was greater than in the discovery cohort, the high median ARI indicates that subtype assignments were generally well preserved under resampling.

Statistical evaluation of canonical Alzheimer’s disease biomarkers demonstrated that the replicated proteomic groups were associated with differences in established Alzheimer’s disease biomarkers. Significant differences were observed in total tau (t-tau; p = 4.32 × 10⁻⁵) and phosphorylated tau (p-tau; p = 1.15 × 10⁻⁵), with higher levels observed in Subtype 1. Amyloid-β42 (Aβ42) levels also differed between groups (p = 5.72 × 10⁻³), as did the Aβ42/Aβ40 ratio (p = 6.80 × 10⁻⁶) (Figure 5). Because proteins were selected based on differences between cognitively normal and AD individuals, these biomarker associations should be interpreted as characterization of the resulting proteomic groups rather than as independent validation of the group structure.

**FIGURE 5.**
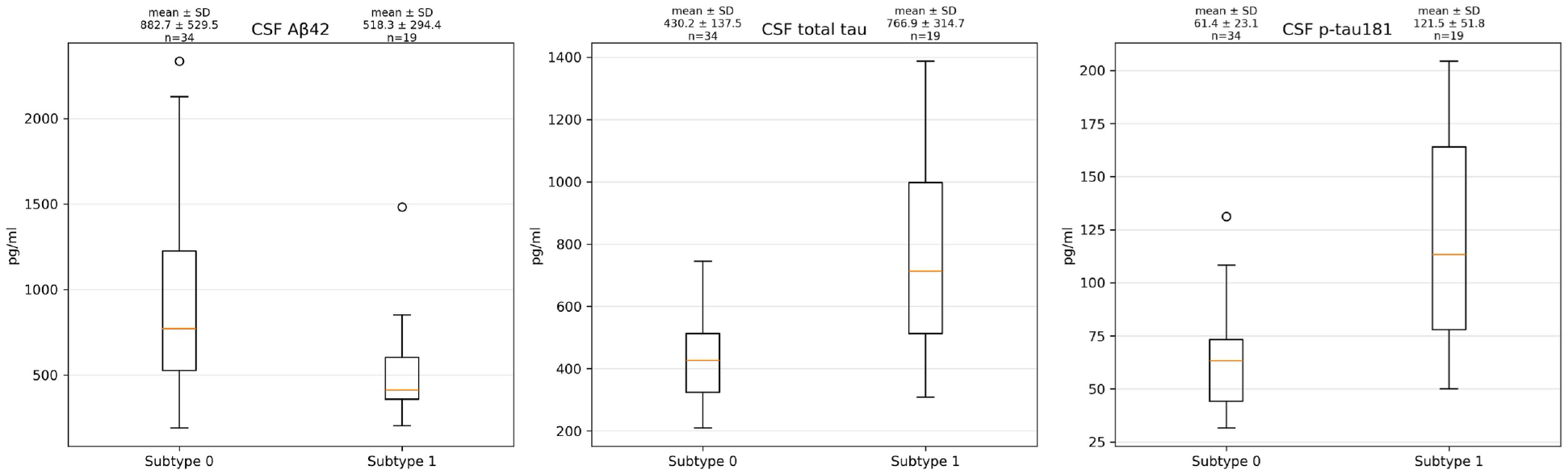
Replication cohort biomarker differences across PH-derived subtypes. Boxplots show significant differences between Subtype 0 and Subtype 1 in Aβ42 (p = 5.72×10⁻³), total tau (p = 4.32×10⁻⁵), and phosphorylated tau (p = 1.15×10⁻⁵). Values above each boxplot show the mean ± SD and sample size. Subtype 1 exhibits elevated tau and p-tau levels and reduced Aβ42, consistent with increased Alzheimer’s disease pathology.

Together, these results demonstrate that a comparable two-group organization can be recovered in an independent cohort profiled using a different proteomic platform and that these groups are associated with differences in core Alzheimer’s disease biomarkers.

### 3.8 | Clinical and biomarker characteristics

To characterize the biological and clinical profiles of the identified subtypes, we compared demographic variables, cognitive measures, and CSF biomarkers across groups.

Summary visualizations of subtype-level biomarker, clinical, and genetic characteristics are provided in Figure S2. In the original cohort (Table 1), unadjusted group comparisons showed that subtype 1 exhibited substantially higher levels of total tau and phosphorylated tau compared to subtype 0 (total tau: 460.5 ± 101.7 vs 283.1 ± 94.7; p-tau: 47.4 ± 11.6 vs 27.6 ± 10.2). Aβ42 levels also differed between subtypes (659.7 ± 251.0 vs 600.3 ± 246.1; p = 0.030), although this difference was considerably smaller than the differences observed for total tau and p-tau. Thus, subtype differentiation in the discovery cohort was predominantly characterized by tau-related pathology, with a more modest difference in Aβ42. Cognitive performance, as measured by MMSE, was comparable between subtypes (23.5 ± 2.1 vs 23.3 ± 2.0), and age distributions were also similar. Notably, the proportion of APOE ε4 carriers was higher in subtype 1 than in subtype 0 (78.6% vs 63.4%).

**TABLE 1.** Clinical and cerebrospinal fluid biomarker characteristics of the identified dominant proteomic subtypes in the discovery cohort.

| **Characteristic<sup>a</sup>** | **Controls n = 167** | **Subtype 0 n = 71** | **Subtype 1 n = 42** | **Statistics<sup>b</sup>** |
|---|---|---|---|---|
| Age, years | 74.42 (5.94) | 75.60 (8.49) | 74.06 (9.13) | one-way ANOVA: F = 0.85 (p = 4.29e-01) |
| Female, n (%) | 78 (46.7) | 34 (47.9) | 14 (33.3) | χ² test: χ² = 2.75 (p = 2.53e-01) |
| No. of APOE ε4 alleles (0/1/2) | 126/38/3 | 26/33/12 | 9/22/11 | χ² test: χ² = 65.60 (p = 1.92e-13) |
| APOE ε4 carriers (%) | 24.6 | 63.4 | 78.6 | — |
| MMSE | 29.03 (1.16) | 23.31 (2.03) | 23.55 (2.07) | one-way ANOVA: F = 430.33 (p = 1.06e-85) |
| CSF Aβ42, pg/ml | 1045.19 (368.57) | 600.26 (246.13) | 659.71 (251.00) | one-way ANOVA: F = 51.51 (p = 3.69e-19) |
| CSF p-tau181, pg/ml | 21.75 (8.79) | 27.62 (10.16) | 47.39 (11.60) | one-way ANOVA: F = 119.28 (p = 5.46e-38) |
| CSF total tau, pg/ml | 236.78 (85.86) | 283.09 (94.71) | 460.53 (101.68) | one-way ANOVA: F = 102.14 (p = 6.73e-34) |

In the replication cohort (Table 2), unadjusted group comparisons showed that the tau-related pattern was reproduced, whereas the direction of the Aβ42 difference differed from that observed in the discovery cohort. Subtype 1 demonstrated markedly elevated total tau (766.9 ± 314.7 vs 430.2 ± 137.5) and p-tau levels (121.5 ± 51.8 vs 61.4 ± 23.1). However, unlike in the discovery cohort, Subtype 1 exhibited substantially lower Aβ42 concentrations (518.3 ± 294.4 vs 882.7 ± 529.5). Thus, the tau-associated distinction between subtypes was directionally consistent across cohorts, whereas the amyloid-associated difference was not. These findings indicate a more pronounced AD-like biomarker profile in subtype 1 in the replication cohort, characterized by both increased tau pathology and reduced amyloid levels. Cognitive scores (MoCA) were similar across subtypes (22.6 ± 3.2 vs 22.5 ± 3.8), and no significant differences were observed in age or sex distribution. APOE ε4 carrier frequencies were comparable between the two replication-cohort subtypes (47.4% in Subtype 1 vs 44.1% in Subtype 0; OR = 1.14, Fisher’s exact p = 1.00).

**TABLE 2.** Clinical and cerebrospinal fluid biomarker characteristics of the identified dominant proteomic subtypes in the replication cohort.

| Characteristic<sup>a</sup> | Controls n = 40 | Subtype 0 n = 34 | Subtype 1 n = 19 | Statistics<sup>b</sup> |
|---|---|---|---|---|
| Age, years | 68.88 (6.24) | 74.00 (8.13) | 70.11 (10.62) | one-way ANOVA: F = 3.94 (p = 0.023) |
| Female, n (%) | 21 (52.5) | 17 (50.0) | 11 (57.9) | χ² test: χ² = 0.31 (p = 0.858) |
| No. of APOE ε4 alleles (0/1/2) | 32/8/0 | 19/15/0 | 10/9/0 | χ² test: χ² = 6.51 (p = 0.039) |
| APOE ε4 carriers (%) | 20.0 | 44.1 | 47.4 | — |
| MoCA | 26.57 (2.84) | 22.50 (3.82) | 22.56 (3.24) | one-way ANOVA: F = 14.01 (p = 7.42e-06) |
| CSF Aβ42, pg/ml | 1095.35 (335.45) | 882.68 (529.52) | 518.26 (294.43) | one-way ANOVA: F = 12.74 (p = 1.34e-05) |
| CSF Aβ40, pg/ml | 11155.80 (2820.63) | 11657.76 (4725.19) | 11954.68 (5155.69) | one-way ANOVA: F = 0.28 (p = 0.756) |
| CSF Aβ42/Aβ40 ratio | 0.10 (0.01) | 0.07 (0.02) | 0.04 (0.01) | one-way ANOVA: F = 66.10 (p = 2.17e-18) |
| CSF p-tau181, pg/ml | 30.51 (6.70) | 61.39 (23.10) | 121.48 (51.76) | one-way ANOVA: F = 71.01 (p = 3.11e-19) |
| CSF total tau, pg/ml | 260.70 (211.54) | 430.18 (137.47) | 766.89 (314.71) | one-way ANOVA: F = 35.78 (p = 3.67e-12) |

**Abbreviations:** APOE, apolipoprotein E; CSF, cerebrospinal fluid; Aβ42, amyloid-beta 42; Aβ40, amyloid-beta 40; p-tau181, phosphorylated tau at threonine 181; MMSE, Mini-Mental State Examination; MoCA, Montreal Cognitive Assessment; CDRSB, Clinical Dementia Rating Sum of Boxes; AD, Alzheimer’s disease; ANOVA, analysis of variance. Continuous variables are reported as mean (standard deviation), and categorical variables as counts (percentages). Group differences were assessed using one-way ANOVA for continuous variables and chi-squared (χ²) tests for categorical variables.
Values are presented as mean ± standard deviation or percentage, as appropriate. In both cohorts, subtype 1 exhibited higher levels of total tau and phosphorylated tau compared to subtype 0, with consistent trends across datasets. Aβ42 levels were lower in subtype 1 in the replication cohort, while they showed a smaller difference in the original cohort. Cognitive scores (MMSE in the original cohort and MoCA in the replication cohort) were similar between subtypes. APOE ε4 carrier frequency was higher in Subtype 1 in the discovery cohort but comparable between states in the replication cohort.

Taken together, these results demonstrate that Subtype 1 is consistently associated with elevated tau pathology across cohorts, whereas the direction of the Aβ42 difference was cohort-dependent.

### 3.9 | Exploratory substructure within the larger proteomic state

To explore potential heterogeneity within the larger proteomic group, we performed a secondary, exploratory clustering analysis among its 71 individuals. This analysis identified three groups containing 49, 11, and 11 individuals, respectively. Given the small sizes of two of these groups and the absence of an equivalent analysis in the replication cohort, we consider these groups exploratory and hypothesis-generating rather than validated patient subtypes.

The three exploratory groups showed differences in several molecular and demographic characteristics (Table 3). Group 2 (n = 11) had higher mean tau (414.67 ± 117.55 pg/ml) and phosphorylated tau (41.39 ± 12.09 pg/ml) and the highest proportion of APOE ε4 carriers (81.8%). Group 1 (n = 11) had comparatively higher mean CSF Aβ42 levels (887.65 ± 414.94 pg/ml) and was the oldest group (mean age 82.71 ± 7.56 years), whereas the largest group (n = 49) had comparatively lower mean tau and phosphorylated tau levels (252.51 ± 59.49 pg/ml and 24.68 ± 6.62 pg/ml, respectively). These profiles indicate molecular heterogeneity within the larger proteomic group but, given the small sample sizes, are not interpreted as evidence for distinct disease mechanisms or trajectories.

**TABLE 3.**
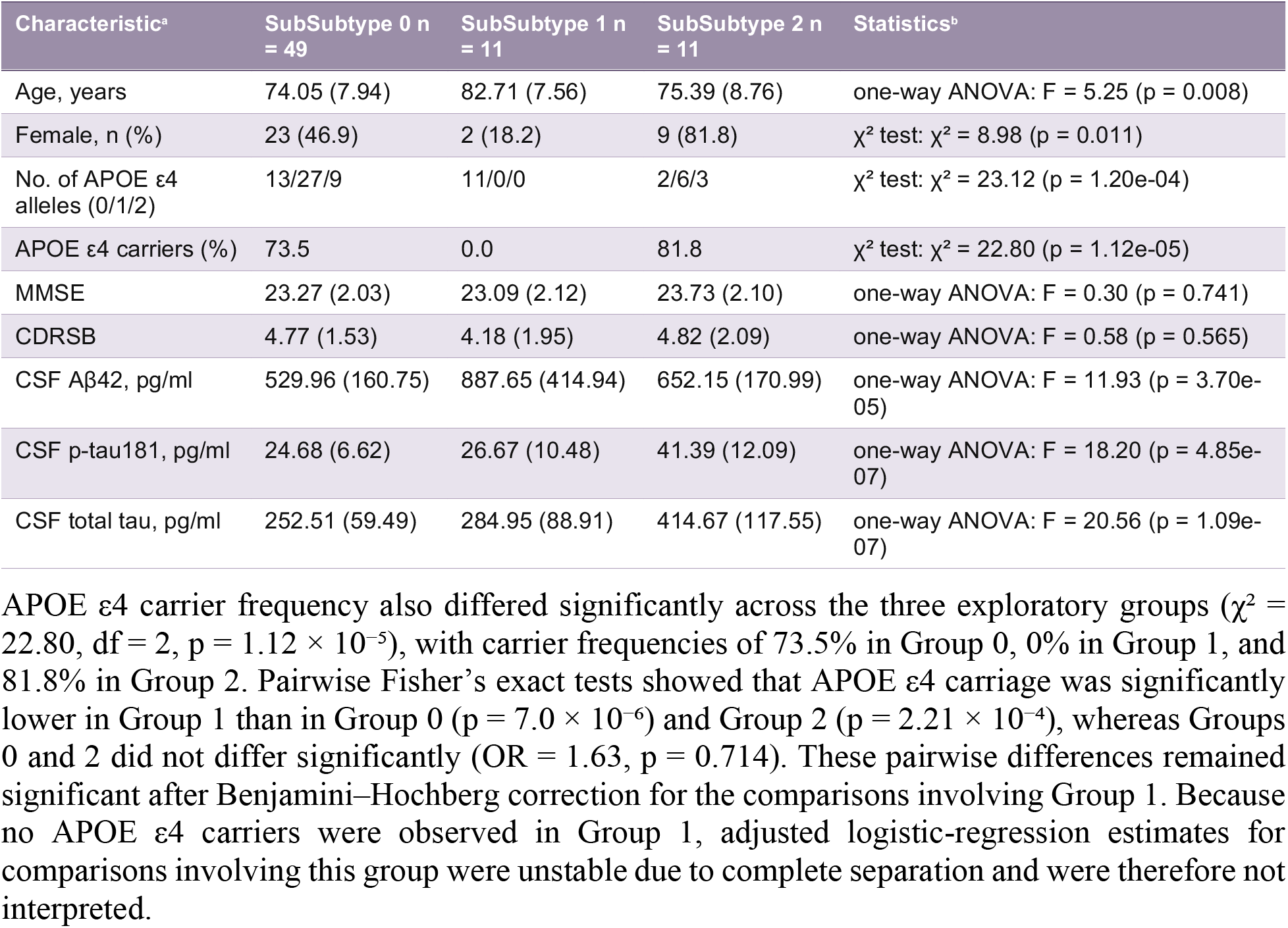
Demographic, genetic, cognitive, and cerebrospinal fluid biomarker characteristics of the sub-subtypes identified within the largest dominant proteomic subgroup in the discovery cohort.

| Characteristic<sup>a</sup> | SubSubtype 0 n = 49 | SubSubtype 1 n = 11 | SubSubtype 2 n = 11 | Statistics<sup>b</sup> |
|---|---|---|---|---|
| Age, years | 74.05 (7.94) | 82.71 (7.56) | 75.39 (8.76) | one-way ANOVA: F = 5.25 (p = 0.008) |
| Female, n (%) | 23 (46.9) | 2 (18.2) | 9 (81.8) | χ² test: χ² = 8.98 (p = 0.011) |
| No. of APOE ε4 alleles (0/1/2) | 13/27/9 | 11/0/0 | 2/6/3 | χ² test: χ² = 23.12 (p = 1.20e-04) |
| APOE ε4 carriers (%) | 73.5 | 0.0 | 81.8 | χ² test: χ² = 22.80 (p = 1.12e-05) |
| MMSE | 23.27 (2.03) | 23.09 (2.12) | 23.73 (2.10) | one-way ANOVA: F = 0.30 (p = 0.741) |
| CDRSB | 4.77 (1.53) | 4.18 (1.95) | 4.82 (2.09) | one-way ANOVA: F = 0.58 (p = 0.565) |
| CSF Aβ42, pg/ml | 529.96 (160.75) | 887.65 (414.94) | 652.15 (170.99) | one-way ANOVA: F = 11.93 (p = 3.70e-05) |
| CSF p-tau181, pg/ml | 24.68 (6.62) | 26.67 (10.48) | 41.39 (12.09) | one-way ANOVA: F = 18.20 (p = 4.85e-07) |
| CSF total tau, pg/ml | 252.51 (59.49) | 284.95 (88.91) | 414.67 (117.55) | one-way ANOVA: F = 20.56 (p = 1.09e-07) |

APOE ε4 carrier frequency also differed significantly across the three exploratory groups (χ² = 22.80, df = 2, p = 1.12 × 10⁻⁵), with carrier frequencies of 73.5% in Group 0, 0% in Group 1, and 81.8% in Group 2. Pairwise Fisher’s exact tests showed that APOE ε4 carriage was significantly lower in Group 1 than in Group 0 (p = 7.0 × 10⁻⁶) and Group 2 (p = 2.21 × 10⁻⁴), whereas Groups 0 and 2 did not differ significantly (OR = 1.63, p = 0.714). These pairwise differences remained significant after Benjamini–Hochberg correction for the comparisons involving Group 1. Because no APOE ε4 carriers were observed in Group 1, adjusted logistic-regression estimates for comparisons involving this group were unstable due to complete separation and were therefore not interpreted.

Cognitive measures were comparatively similar across the three groups, with mean MMSE scores of approximately 23 in each group (Table 3), despite the observed differences in molecular and demographic characteristics. Differences in sex distribution were also observed across the groups. Together, these exploratory findings suggest that the larger proteomic group may contain additional internal heterogeneity associated with CSF biomarker, APOE ε4, age, and sex profiles. However, the two smaller groups each contain only 11 individuals, and this finer-scale structure was not evaluated in the independent replication cohort. These findings should therefore be considered hypothesis-generating and will require confirmation in larger independent cohorts before they can be interpreted as reproducible biological subtypes or distinct disease trajectories.

## 4 | DISCUSSION

In this study, we used diffusion geometry and persistent homology to characterize the multiscale and partially continuous organization of the Alzheimer’s proteomic landscape. H₀ identified two dominant, reproducible groups at a coarse scale, while H₁ revealed continuous and nonlinear relationships beyond discrete partitions. The two states were further characterized by differences in tau and amyloid biomarker profiles and by substantial enrichment of APOE ε4 carriage relative to controls, consistent with the established biological context of Alzheimer’s disease described by the NIA-AA research framework [1] and AD GWAS [13].

A central feature of this approach is its unsupervised nature: proteomic state structure was inferred directly from the geometry of the proteomic data, without using diagnostic labels or imposing a predefined number of groups [32]. Examination of the H₀ persistence spectrum revealed two prominent connected components at a coarse filtration scale, representing one level of the multiscale connectivity hierarchy [20, 33]. Beyond connected components, persistent H₁ features identified loop-like structures, indicating continuous, nonlinear organization beyond the H₀ structure. Importantly, comparison with Gaussian null data demonstrated that the observed H₁ persistence was unlikely to arise from unstructured high-dimensional variation alone, providing statistical support for the presence of non-random higher-order topology in the proteomic landscape. This interpretation is consistent with the hierarchical substructure observed within the dominant state, where three distinct subgroups exhibited divergent biomarker profiles despite belonging to the same large-scale connected component.

Across both discovery and replication cohorts, the most consistent molecular differences between proteomic states were observed in tau-related biomarkers. The high-tau subtype exhibited substantially elevated levels of total tau and phosphorylated tau, while Aβ42 displayed an opposing pattern, particularly in the replication cohort. These findings align with established biomarker relationships in Alzheimer’s disease [4, 15].

Notably, cognitive measures and demographic variables did not differ significantly between groups, suggesting that the identified molecular differences are not accompanied by clear cross-sectional differences in clinical severity. Although this pattern may be consistent with distinct molecular states rather than sequential disease stages, the cross-sectional design does not allow their temporal relationships or trajectories to be established. Longitudinal follow-up will therefore be necessary to determine whether these subtypes represent stable molecular phenotypes or different stages of disease progression.

Recent large-scale CSF proteomic studies have demonstrated substantial molecular heterogeneity in Alzheimer’s disease. Notably, Tijms et al. identified five molecular subtypes characterized by distinct biological pathways and genetic risk profiles [18]. Our findings complement this work by providing a different perspective on how such heterogeneity is organized. Rather than representing the proteomic landscape solely through discrete subtype assignments, our topology-based framework reveals molecular organization across multiple scales, with broad proteomic states accompanied by finer-scale substructure and continuous higher-order relationships. Persistent homology is particularly suited to this setting because it does not require the proteomic landscape to be represented by a single clustering solution or at a single resolution; instead, it tracks how structure emerges, persists, and changes across filtration scales [25, 32, 34, 35]. From this perspective, previously reported molecular subtypes and the proteomic states identified here need not represent competing descriptions of Alzheimer’s disease heterogeneity, but may reflect complementary levels of organization within a complex and partially continuous molecular landscape. Topology-based approaches have also previously been used for disease subtyping outside Alzheimer’s disease. For example, Nicolau et al. applied topological data analysis to breast cancer transcriptomic data and identified a molecularly and clinically distinct subgroup that was not recovered by conventional clustering methods [36]. In this context, the novelty of the present study lies not in the use of topology for disease stratification itself, but in its application to the multiscale organization of the Alzheimer’s disease CSF proteomic landscape, integrating persistent homology with molecular, biomarker, and genetic characterization.

Notably, the three sub-subtypes identified within the dominant proteomic cluster show partial correspondence with previously reported Alzheimer’s disease subtype classifications derived from supervised machine learning approaches [11]. Together, these findings suggest that disease-related variation may exist as substructure within broader proteomic states, highlighting the ability of persistent homology to capture multiscale organization in high-dimensional proteomic data [33, 25].

In our analysis, a similar three-group structure emerged at a finer level of resolution within the dominant subtype, with distinct biological interpretations. One subgroup exhibited a low-tau profile, whereas a second subgroup showed a pronounced tau-elevated signature. Notably, this latter group may correspond most closely to the tau-elevated subtype reported by Hou et al. [11], although further analysis would be required to confirm direct alignment.

In contrast, a third subgroup displayed relatively elevated CSF Aβ42 alongside only moderate tau pathology, a profile that does not align with a classical amyloid-driven disease trajectory. While this interpretation remains tentative, these findings highlight the heterogeneity captured within the proteomic substructure and underscore the importance of distinguishing between amyloid- and tau-driven mechanisms.

Importantly, while prior studies identified these subtypes as primary groupings, our findings suggest that such disease trajectories may also exist as substructure within broader proteomic states. Persistent homology therefore captures this finer-scale proteomic organization [33, 25].

Beyond this multiscale proteomic organization, we examined whether the identified states were also associated with established genetic risk for AD. APOE ε4 carrier frequency was higher in the high-tau than low-tau state in the discovery cohort (78.6% vs 63.4%), but this difference was not replicated in the independent cohort (47.4% vs 44.1%; OR = 1.14, Fisher’s exact p = 1.00). Thus, although APOE ε4 may contribute to the biological context of the identified states, its differential enrichment was not consistent across cohorts [22].

Proteomic analyses further linked state differences to synaptic and neuronal processes, including synaptic organization, neuronal differentiation, and cell–cell communication [37]. Differential proteins included DLG2 and STMN2, involved in synaptic organization and axonal maintenance, respectively, together with alterations in Eph receptor signaling and 14-3-3 proteins, providing additional links between the proteomic signatures, synaptic dysfunction, and tau-related biology.

A key strength of this study is the replication of results across independent cohorts profiled on fundamentally different proteomic platforms. The discovery cohort used aptamer-based SomaScan profiling, whereas the Johns Hopkins replication cohort used mass spectrometry, providing substantially different measurement modalities.

The recovery of comparable proteomic organization across these modalities indicates that the observed structure is not specific to a single proteomic measurement platform. However, because disease-associated proteins were selected using CN–AD comparisons in both analyses, the accompanying associations with canonical AD biomarkers and APOE ε4 should not be regarded as fully independent biological validation. Rather, these findings demonstrate that the recovered proteomic groups are associated with established axes of AD-related molecular heterogeneity across two distinct proteomic platforms.

Several limitations should be acknowledged. Sample sizes remain modest, and larger datasets will be necessary to confirm the robustness of subtype structure and its association with genetic and clinical variables. Longitudinal studies will be particularly important to determine whether these proteomic states differ in disease progression or treatment response. In addition, the absence of direct measurement of co-pathologies in this cohort represents an important limitation, as Alzheimer’s disease frequently co-exists with other pathological processes, including α-synuclein, TDP-43, and vascular pathology. The presence of such co-morbidities may contribute to additional heterogeneity and potentially give rise to further subtypes not captured in the present analysis. Finally, integration with imaging and genomic modalities may provide a more comprehensive view of the biological axes identified here.

An additional limitation arises from the disease-informed feature-selection strategy. Proteins entering the topological analysis were pre-selected based on differences between cognitively normal and AD individuals. This enrichment for disease-associated proteins may predispose the resulting proteomic organization to align with established AD-related molecular axes, including amyloid and tau pathology and their genetic correlates. Consequently, the observed associations with CSF biomarkers and APOE ε4 should not be interpreted as fully independent validation of the identified groups. Rather, they characterize the biological variation captured by a feature space intentionally enriched for AD-associated proteins. Future studies using disease-agnostic feature selection, independent protein sets, or analyses excluding proteins closely associated with canonical AD biomarker pathways will be important for determining the extent to which the observed organization persists independently of this selection strategy.

In conclusion, this study identifies a reproducible large-scale organization of the Alzheimer’s disease proteomic landscape across independent cohorts and orthogonal proteomic platforms. The H₀ analysis identifies two dominant proteomic states within this organization, while higher-dimensional persistent homology provides evidence for nontrivial cyclic structure that is not fully captured by discrete partitions. The proteomic states are associated with differences in tau pathology, amyloid biomarkers, and APOE ε4 carrier frequency, although these associations should be interpreted in the context of the disease-informed feature-selection strategy. The convergence of topological, proteomic, biomarker, and genetic evidence supports a coherent biological framework and highlights the potential of topology-based methods to refine disease stratification and inform more targeted therapeutic strategies.

## DATA AVAILABILITY STATEMENT

The data used in this study were obtained from the ADNI database (adni.loni.usc.edu) and the Johns Hopkins’s database. Researchers can apply for ADNI data through the ADNI website. Detailed information on data collection procedures are found on the portal.

## CODE AVAILABILITY STATEMENT

All software used in this study is publicly available. The code used in this study can be accessed at https://github.com/KellyShahu/Proteomic_analysis_TDA_Alzheimer-s

## ACKNOWLEDGEMENTS

Data collection and sharing for the Alzheimer’s Disease Neuroimaging Initiative (ADNI) is funded by the National Institute on Aging (National Institutes of Health Grant U19 AG024904). The grantee organization is the Northern California Institute for Research and Education.

In the past, ADNI has also received funding from the National Institute of Biomedical Imaging and Bioengineering, the Canadian Institutes of Health Research, and private sector contributions through the Foundation for the National Institutes of Health (FNIH) including generous contributions from the following: AbbVie, Alzheimer’s Association; Alzheimer’s Drug Discovery Foundation; Araclon Biotech; BioClinica, Inc.; Biogen; Bristol-Myers Squibb Company; CereSpir, Inc.; Cogstate; Eisai Inc.; Elan Pharmaceuticals, Inc.; Eli Lilly and Company; EuroImmun; F. Hoffmann-La Roche Ltd and its affiliated company Genentech, Inc.; Fujirebio; GE Healthcare; IXICO Ltd.; Janssen Alzheimer Immunotherapy Research & Development, LLC.; Johnson & Johnson Pharmaceutical Research &Development LLC.; Lumosity; Lundbeck; Merck & Co., Inc.; Meso Scale Diagnostics, LLC.; NeuroRx Research; Neurotrack Technologies; Novartis Pharmaceuticals Corporation; Pfizer Inc.; Piramal Imaging; Servier; Takeda Pharmaceutical Company; and Transition Therapeutics.

## CONFLICT OF INTEREST STATEMENT

The authors declare no competing interests.

## Funding information

Duke Science and Technology LAUNCH; Bass

Connections at Duke University; National

Institute on Aging, National Institutes of

Health (NIA, NIH), Grant/Award Number:

P30AG07295

## CONSENT STATEMENT

The Alzheimer’s Disease Neuroimaging Initiative (ADNI) study was approved by the Institutional Review Boards (IRBs) of all participating sites. Written informed consent was obtained from all participants or their authorized representatives prior to their inclusion in the study.

For the replication cohort from Johns Hopkins, all participants signed institutional review board– approved consent forms before participating in the study.

## SUPPLEMENTARY MATERIAL

**FIGURE S1.**
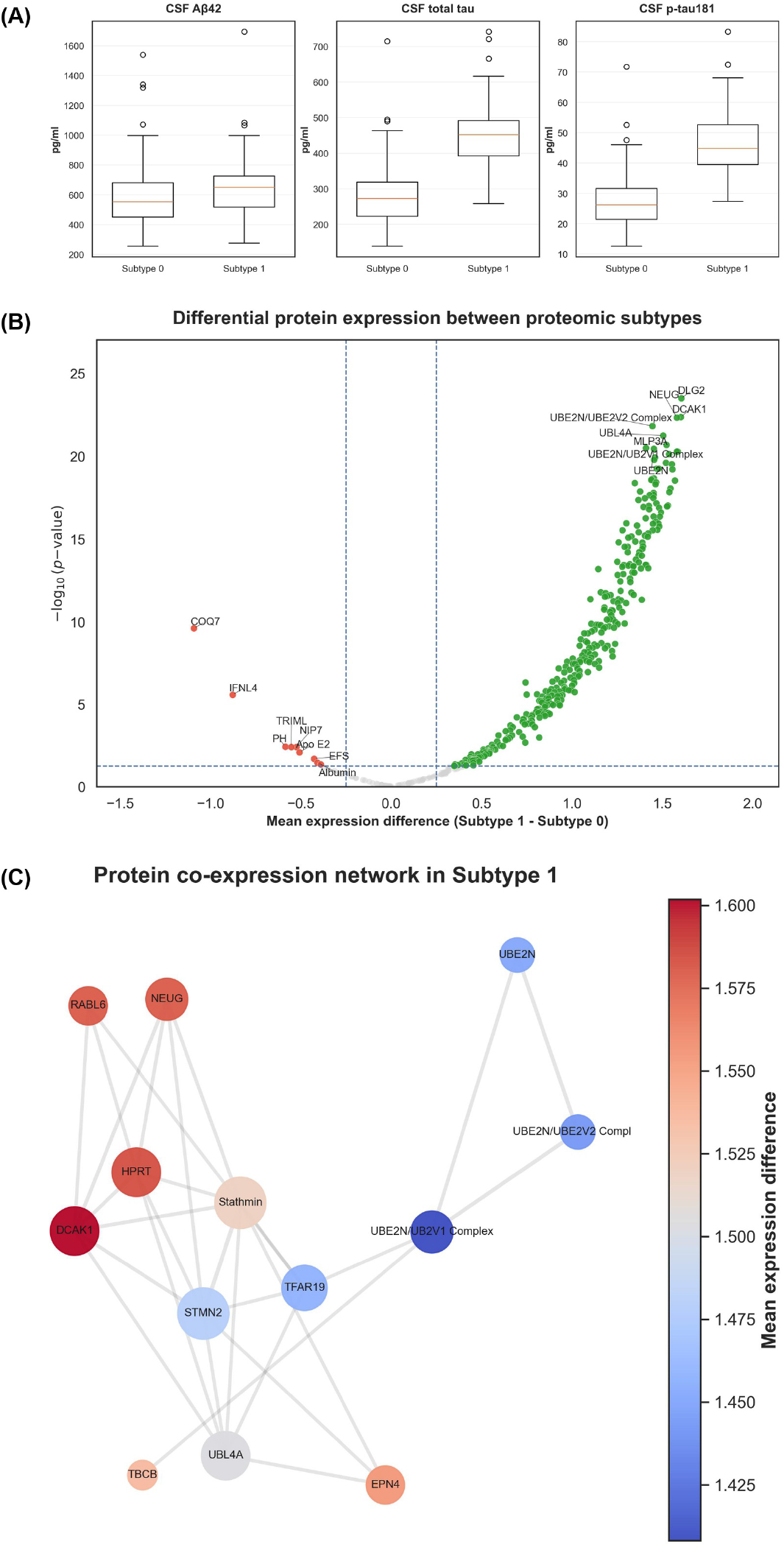
Molecular and proteomic differences between dominant Alzheimer’s disease proteomic subtypes. **(A)** Differences in cerebrospinal fluid (CSF) biomarkers across proteomic subtypes. Boxplots show the distributions of Aβ42, total tau (TAU), and phosphorylated tau (PTAU) across the two dominant subtypes. Subtype 1 exhibits elevated TAU and PTAU levels, together with a smaller but statistically significant difference in Aβ42, consistent with distinct molecular disease profiles between subtypes. Values above each boxplot represent the mean ± standard deviation (SD) for each group, with sample size (n) indicated. **(B)** Differential protein expression between proteomic subtypes. Volcano plot displaying mean differences in protein abundance between Subtype 1 and Subtype 0 (x-axis) against statistical significance (−log₁₀ p-value, y-axis). Each point represents a protein. Proteins passing the significance threshold (false discovery rate < 0.05) and effect size cutoff are highlighted, with positive values indicating increased expression in Subtype 1 and negative values indicating increased expression in Subtype 0. Selected top-associated proteins are labeled. The distribution demonstrates a marked asymmetry, with a substantially larger set of proteins upregulated in Subtype 1. **(C)** Protein co-expression network associated with Subtype 1. Nodes represent subtype-associated proteins and edges indicate significant pairwise correlations above a defined threshold. Node size reflects network connectivity (degree), while node color represents the mean expression difference between Subtype 1 and Subtype 0. The resulting network reveals a densely interconnected module enriched for proteins involved in synaptic signaling and cellular communication, suggesting coordinated molecular regulation within Subtype 1.

**FIGURE S2.**
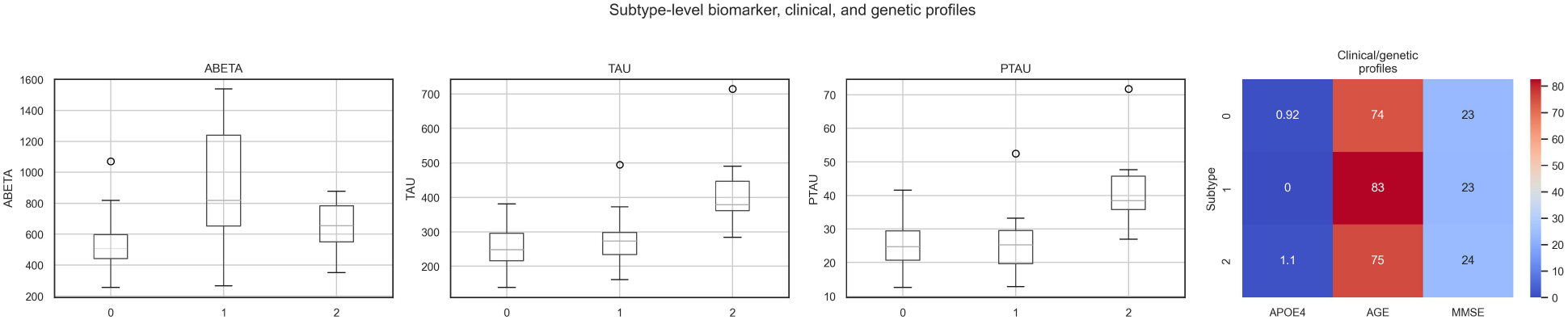
Biomarker and clinical characterization of PH-derived sub-subtypes. Distribution of core CSF biomarkers (Aβ42, total tau, and p-tau) across sub-subtypes identified within the dominant proteomic subgroup. Boxplots illustrate median, interquartile range, and outliers. Sub-subtypes exhibit distinct biomarker profiles, with differential patterns in amyloid and tau pathology, supporting biological differentiation beyond the primary subtype level. Heatmap summarizing subtype-level clinical and genetic features, including APOE ε4 carrier frequency, mean age, and MMSE scores. Color intensity reflects relative magnitude across sub-subtypes. The observed variation highlights differences in genetic risk and clinical presentation, further supporting the presence of meaningful substructure within the dominant proteomic subtype.

**TABLE S1.** Pathway enrichment analysis of proteomic subtypes derived from persistent homology. Significant pathways identified through gene ontology and pathway enrichment analysis across Alzheimer’s disease subtypes discovered via persistent homology (PH). Enrichment results span biological process, cellular component, and molecular function categories, highlighting subtype-specific pathway activation patterns. Subtypes exhibit distinct enrichment profiles, with differential representation of synaptic, neuronal, and signaling pathways. The observed pathway heterogeneity supports the biological validity of PH-derived subtype stratification and suggests divergent molecular mechanisms underlying disease progression across subtypes.

| source | native | name | p_value | term_size | intersection_size | query_size |
|---|---|---|---|---|---|---|
| GO:CC | GO:0045202 | synapse | 2.38e-09 | 1608 | 31 | 105 |
| GO:CC | GO:0030054 | cell junction | 1.23e-07 | 2370 | 35 | 105 |
| GO:BP | GO:0023052 | signaling | 1.32e-07 | 6558 | 63 | 100 |
| GO:BP | GO:0007154 | cell communication | 1.76e-07 | 6598 | 63 | 100 |
| GO:BP | GO:0099536 | synaptic signaling | 2.35e-06 | 803 | 20 | 100 |
| GO:BP | GO:0007267 | cell-cell signaling | 3.83e-06 | 1316 | 25 | 100 |
| GO:BP | GO:0007268 | chemical synaptic transmission | 6.84e-06 | 765 | 19 | 100 |
| GO:BP | GO:0098916 | anterograde trans-synaptic signaling | 6.84e-06 | 765 | 19 | 100 |
| GO:BP | GO:0099537 | trans-synaptic signaling | 7.94e-06 | 772 | 19 | 100 |
| GO:BP | GO:0050808 | synapse organization | 2.04e-05 | 561 | 16 | 100 |
| GO:CC | GO:0005829 | cytosol | 2.50e-05 | 5607 | 52 | 105 |
| GO:CC | GO:0005737 | cytoplasm | 4.97e-05 | 12601 | 85 | 105 |
| GO:CC | GO:0098793 | presynapse | 6.05e-05 | 610 | 15 | 105 |
| REAC | REAC:R-HSA-8875656 | MET receptor recycling | 7.62e-05 | 9 | 4 | 70 |
| KEGG | KEGG:04360 | Axon guidance | 1.32e-04 | 183 | 9 | 54 |
| GO:BP | GO:0065008 | regulation of biological quality | 2.78e-04 | 2963 | 35 | 100 |
| GO:MF | GO:0005515 | protein binding | 3.13e-04 | 15082 | 99 | 106 |
| GO:CC | GO:0098839 | postsynaptic density membrane | 4.55e-04 | 117 | 7 | 105 |
| GO:BP | GO:0023051 | regulation of signaling | 4.65e-04 | 3473 | 38 | 100 |
| GO:BP | GO:0010646 | regulation of cell communication | 4.91e-04 | 3480 | 38 | 100 |
| GO:CC | GO:0045211 | postsynaptic membrane | 5.98e-04 | 304 | 10 | 105 |
| GO:CC | GO:0098978 | glutamatergic synapse | 6.31e-04 | 548 | 13 | 105 |
| GO:CC | GO:0098794 | postsynapse | 6.50e-04 | 737 | 15 | 105 |
| GO:CC | GO:0098984 | neuron to neuron synapse | 7.37e-04 | 387 | 11 | 105 |
| GO:BP | GO:0034330 | cell junction organization | 7.94e-04 | 829 | 17 | 100 |
| HPA | HPA:0100211 | Cerebral cortex; neuropil[≥Low] | 1.10e-03 | 4999 | 57 | 81 |
| GO:BP | GO:0051128 | regulation of cellular component organization | 1.21e-03 | 2427 | 30 | 100 |
| GO:BP | GO:0007416 | synapse assembly | 1.26e-03 | 266 | 10 | 100 |
| GO:CC | GO:0014069 | postsynaptic density | 1.49e-03 | 337 | 10 | 105 |
| GO:BP | GO:0007399 | nervous system development | 1.57e-03 | 2597 | 31 | 100 |
| GO:CC | GO:0099634 | postsynaptic specialization membrane | 1.73e-03 | 143 | 7 | 105 |
| MIRNA | MIRNA:hsa-mir-338-5p | hsa-mir-338-5p | 1.83e-03 | 1349 | 24 | 104 |
| GO:CC | GO:0097060 | synaptic membrane | 1.84e-03 | 426 | 11 | 105 |
| GO:CC | GO:0032279 | asymmetric synapse | 2.19e-03 | 352 | 10 | 105 |
| HPA | HPA:0100212 | Cerebral cortex; neuropil[≥Medium] | 2.91e-03 | 2616 | 37 | 81 |
| GO:BP | GO:0072657 | protein localization to membrane | 3.04e-03 | 445 | 12 | 100 |
| GO:CC | GO:0099572 | postsynaptic specialization | 3.38e-03 | 370 | 10 | 105 |
| REAC | REAC:R-HSA-428542 | Regulation of commissural axon pathfinding by SLIT and ROBO | 8.35e-03 | 9 | 3 | 70 |
| GO:CC | GO:0005576 | extracellular region | 1.03e-02 | 4249 | 38 | 105 |
| GO:BP | GO:0050804 | modulation of chemical synaptic transmission | 1.07e-02 | 503 | 12 | 100 |
| HPA | HPA:0250142 | Hippocampus; neuronal projections[≥Medium] | 1.08e-02 | 10 | 3 | 81 |
| GO:BP | GO:0099177 | regulation of trans-synaptic signaling | 1.09e-02 | 504 | 12 | 100 |
| GO:CC | GO:0031594 | neuromuscular junction | 1.12e-02 | 78 | 5 | 105 |
| GO:BP | GO:0032956 | regulation of actin cytoskeleton organization | 1.14e-02 | 340 | 10 | 100 |
| GO:BP | GO:0031175 | neuron projection development | 1.17e-02 | 1010 | 17 | 100 |
| GO:BP | GO:0021954 | central nervous system neuron development | 1.27e-02 | 94 | 6 | 100 |
| MIRNA | MIRNA:hsa-mir-28-3p | hsa-mir-28-3p | 1.28e-02 | 504 | 13 | 104 |
| GO:BP | GO:0007275 | multicellular organism development | 1.89e-02 | 4690 | 42 | 100 |
| GO:CC | GO:0031982 | vesicle | 1.92e-02 | 4032 | 36 | 105 |
| HPA | HPA:0250141 | Hippocampus; neuronal projections[≥Low] | 1.95e-02 | 12 | 3 | 81 |
| GO:BP | GO:0048666 | neuron development | 1.99e-02 | 1167 | 18 | 100 |
| GO:CC | GO:0005615 | extracellular space | 2.32e-02 | 3252 | 31 | 105 |
| HPA | HPA:0100152 | Cerebral cortex; synapses[≥Medium] | 2.53e-02 | 13 | 3 | 81 |
| TF | TF:M08568_1 | Factor: ER71:E2A; motif: CASSTGNACCGGAWRYN; match class: 1 | 2.87e-02 | 15344 | 99 | 107 |
| GO:BP | GO:0030182 | neuron differentiation | 2.88e-02 | 1444 | 20 | 100 |
| TF | TF:M00428_1 | Factor: E2F-1; motif: NKTSSCGC; match class: 1 | 2.98e-02 | 3602 | 38 | 107 |
| GO:CC | GO:0098685 | Schaffer collateral - CA1 synapse | 3.02e-02 | 96 | 5 | 105 |
| TF | TF:M13142_1 | Factor: ZBTB33; motif: GGYGCTG; match class: 1 | 3.03e-02 | 3317 | 36 | 107 |
| GO:BP | GO:0065007 | biological regulation | 3.09e-02 | 12804 | 81 | 100 |
| HPA | HPA:0090991 | Cerebellum; molecular layer - neuropil[≥Low] | 3.12e-02 | 263 | 9 | 81 |
| GO:BP | GO:0032970 | regulation of actin filament-based process | 3.14e-02 | 382 | 10 | 100 |
| GO:MF | GO:0061650 | ubiquitin-like protein conjugating enzyme activity | 3.45e-02 | 40 | 4 | 106 |
| GO:BP | GO:0099173 | postsynapse organization | 3.46e-02 | 233 | 8 | 100 |
| GO:MF | GO:0035591 | signaling adaptor activity | 3.56e-02 | 128 | 6 | 106 |
| GO:MF | GO:0005488 | binding | 3.77e-02 | 18447 | 106 | 106 |
| GO:BP | GO:0044087 | regulation of cellular component biogenesis | 4.60e-02 | 1006 | 16 | 100 |
| HPA | HPA:0100151 | Cerebral cortex; synapses[≥Low] | 4.87e-02 | 16 | 3 | 81 |
| GO:BP | GO:0035865 | cellular response to potassium ion | 4.88e-02 | 12 | 3 | 100 |
| GO:BP | GO:0035685 | helper T cell diapedesis | 4.89e-02 | 2 | 2 | 100 |
| CORUM | CORUM:2476 | CRKL-PDGFRA-CRK-RAPGEF1 complex | 4.99e-02 | 3 | 2 | 22 |

**TABLE S2.** Differential protein expression between proteomic Subtype 0 and Subtype 1. Positive mean differences indicate higher expression in Subtype 1. FDR correction by Benjamini–Hochberg.

| Protein | Mean (Subtype 0) | Mean (Subtype 1) | Mean Difference (Subtype 1 − 0) | p-value | FDR (BH) |
|---|---|---|---|---|---|
| DLG2 | -0.8729 | 0.8674 | 1.7404 | 2.636e-25 | 1.065e-22 |
| NEUG | -0.8255 | 0.8974 | 1.7229 | 3.020e-24 | 6.100e-22 |
| DCAK1 | -0.8394 | 0.8897 | 1.7292 | 7.092e-24 | 9.551e-22 |
| LRTM2 | -0.7822 | 0.8819 | 1.6641 | 1.367e-23 | 1.381e-21 |
| MLP3A | -0.8263 | 0.8433 | 1.6696 | 2.689e-23 | 2.134e-21 |
| UBL4A | -0.8098 | 0.8262 | 1.636 | 3.169e-23 | 2.134e-21 |
| EphA5 | -0.7767 | 0.8603 | 1.637 | 3.853e-23 | 2.224e-21 |
| HPRT | -0.8544 | 0.8918 | 1.7462 | 1.424e-22 | 7.192e-21 |
| Stathmin | -0.8533 | 0.8544 | 1.7077 | 1.621e-22 | 7.278e-21 |
| STMN2 | -0.8256 | 0.8347 | 1.6603 | 2.832e-22 | 1.144e-20 |
| HAGHL | -0.8157 | 0.8935 | 1.7092 | 9.172e-22 | 3.369e-20 |
| sICAM-5 | -0.7822 | 0.8712 | 1.6533 | 1.110e-21 | 3.736e-20 |
| RABL6 | -0.8147 | 0.8857 | 1.7005 | 1.369e-21 | 4.064e-20 |
| UBE2N/UBE2V2 Complex | -0.7755 | 0.7542 | 1.5297 | 1.408e-21 | 4.064e-20 |
| TFAR19 | -0.7974 | 0.7934 | 1.5908 | 1.581e-21 | 4.258e-20 |
| TBCB | -0.8296 | 0.8132 | 1.6429 | 2.740e-21 | 6.918e-20 |
| VISL1 | -0.7239 | 0.8309 | 1.5548 | 3.702e-21 | 8.797e-20 |
| MLP3B | -0.8029 | 0.7811 | 1.5839 | 5.824e-21 | 1.307e-19 |
| EPN4 | -0.7812 | 0.8982 | 1.6794 | 6.457e-21 | 1.373e-19 |
| SUMO3 | -0.7554 | 0.7303 | 1.4857 | 6.876e-21 | 1.389e-19 |
| UBE2N | -0.7522 | 0.7771 | 1.5293 | 1.095e-20 | 2.107e-19 |
| T132B | -0.7725 | 0.8692 | 1.6417 | 1.624e-20 | 2.982e-19 |
| Thioredoxin reductase 1 | -0.8245 | 0.8754 | 1.7 | 2.093e-20 | 3.677e-19 |
| HOP | -0.7896 | 0.9177 | 1.7073 | 2.622e-20 | 4.414e-19 |
| 14-3-3 protein gamma | -0.8032 | 0.8953 | 1.6985 | 2.834e-20 | 4.439e-19 |
| prostatic binding protein | -0.8018 | 0.788 | 1.5898 | 2.920e-20 | 4.439e-19 |
| C1QT4 | -0.7941 | 0.7759 | 1.5701 | 2.966e-20 | 4.439e-19 |
| ROBO2 | -0.7549 | 0.7894 | 1.5443 | 3.483e-20 | 5.026e-19 |
| UBE2N/UB2V1 Complex | -0.7542 | 0.7477 | 1.5019 | 4.081e-20 | 5.678e-19 |
| DCC | -0.7885 | 0.8671 | 1.6556 | 4.216e-20 | 5.678e-19 |
| T132D | -0.7594 | 0.831 | 1.5904 | 7.281e-20 | 9.489e-19 |
| MDHC | -0.7656 | 0.8322 | 1.5978 | 1.118e-19 | 1.411e-18 |
| CBLN4 | -0.7285 | 0.8534 | 1.5819 | 1.301e-19 | 1.593e-18 |
| CRKL | -0.7959 | 0.7147 | 1.5106 | 1.394e-19 | 1.656e-18 |
| SMUF1 | -0.7577 | 0.8071 | 1.5648 | 1.549e-19 | 1.788e-18 |
| ELAV2 | -0.7627 | 0.7803 | 1.543 | 2.451e-19 | 2.750e-18 |
| FAF2 | -0.7955 | 0.8585 | 1.6539 | 4.041e-19 | 4.413e-18 |
| 14-3-3 protein beta/alpha | -0.7745 | 0.803 | 1.5775 | 4.340e-19 | 4.614e-18 |
| 14-3-3 protein zeta/delta | -0.7771 | 0.7993 | 1.5764 | 5.214e-19 | 5.402e-18 |
| NEUM | -0.799 | 0.8742 | 1.6731 | 6.700e-19 | 6.767e-18 |
| H6ST3 | -0.7796 | 0.7813 | 1.5608 | 7.578e-19 | 7.467e-18 |
| NSF1C | -0.7485 | 0.7833 | 1.5319 | 9.233e-19 | 8.882e-18 |
| AMPH | -0.7893 | 0.8164 | 1.6057 | 1.131e-18 | 1.063e-17 |
| NTNG1 | -0.6933 | 0.8893 | 1.5826 | 1.401e-18 | 1.283e-17 |
| BASP | -0.7744 | 0.754 | 1.5284 | 1.437e-18 | 1.283e-17 |
| NETO1 | -0.736 | 0.8471 | 1.5831 | 1.461e-18 | 1.283e-17 |
| PLXA4 | -0.7398 | 0.8808 | 1.6206 | 1.520e-18 | 1.306e-17 |
| SLIT2 | -0.7719 | 0.8533 | 1.6252 | 1.614e-18 | 1.358e-17 |
| PIANP | -0.7646 | 0.7376 | 1.5023 | 2.245e-18 | 1.851e-17 |
| SYT5 | -0.769 | 0.8426 | 1.6116 | 2.374e-18 | 1.918e-17 |
| SH3L2 | -0.7802 | 0.8486 | 1.6289 | 2.865e-18 | 2.270e-17 |
| VWC2 | -0.7735 | 0.7624 | 1.5359 | 2.923e-18 | 2.271e-17 |
| P34 | -0.7236 | 0.7794 | 1.503 | 3.167e-18 | 2.414e-17 |
| HPCL1 | -0.7091 | 0.7176 | 1.4267 | 4.580e-18 | 3.427e-17 |
| Phospholipase C-beta-1 | -0.7862 | 0.8205 | 1.6067 | 6.232e-18 | 4.578e-17 |
| NPTXR | -0.6786 | 0.8772 | 1.5558 | 8.391e-18 | 6.054e-17 |
| EGFL9 | -0.7803 | 0.7858 | 1.5661 | 1.552e-17 | 1.100e-16 |
| PDE1B | -0.74 | 0.8163 | 1.5563 | 2.988e-17 | 2.082e-16 |
| R4RL1 | -0.6611 | 0.8218 | 1.4828 | 3.404e-17 | 2.331e-16 |
| Calcineurin B a | -0.7172 | 0.8619 | 1.5792 | 4.063e-17 | 2.736e-16 |
| TCTP | -0.7823 | 0.8066 | 1.5889 | 4.386e-17 | 2.905e-16 |
| SHAN3 | -0.762 | 0.7879 | 1.5499 | 5.609e-17 | 3.655e-16 |
| MARE3 | -0.7594 | 0.8 | 1.5595 | 5.904e-17 | 3.786e-16 |
| NEGR1 | -0.7038 | 0.8183 | 1.5221 | 8.758e-17 | 5.455e-16 |
| GAS7 | -0.7208 | 0.7119 | 1.4327 | 8.777e-17 | 5.455e-16 |
| Peroxiredoxin-3 | -0.7229 | 0.7822 | 1.5051 | 9.675e-17 | 5.923e-16 |
| IPP2 | -0.7371 | 0.8183 | 1.5554 | 1.162e-16 | 7.007e-16 |
| UNC5H4 | -0.6745 | 0.6387 | 1.3133 | 1.311e-16 | 7.674e-16 |
| SLIK1 | -0.7175 | 0.7729 | 1.4904 | 1.313e-16 | 7.674e-16 |
| NPTX2 | -0.6826 | 0.8063 | 1.4889 | 1.330e-16 | 7.674e-16 |
| UBE2Z | -0.7453 | 0.7643 | 1.5096 | 1.970e-16 | 1.121e-15 |
| UFC1 | -0.7129 | 0.6608 | 1.3736 | 2.166e-16 | 1.216e-15 |
| RGMB | -0.7538 | 0.7737 | 1.5276 | 2.488e-16 | 1.377e-15 |
| CRK | -0.7401 | 0.6332 | 1.3733 | 2.926e-16 | 1.597e-15 |
| SMOC1 | -0.7454 | 0.8336 | 1.579 | 4.726e-16 | 2.531e-15 |
| OX2G | -0.6464 | 0.8447 | 1.4911 | 4.761e-16 | 2.531e-15 |
| GAG2A | -0.7123 | 0.7425 | 1.4548 | 5.641e-16 | 2.960e-15 |
| TMOD3 | -0.7174 | 0.6667 | 1.384 | 6.487e-16 | 3.360e-15 |
| IDI1 | -0.708 | 0.7731 | 1.4811 | 1.126e-15 | 5.761e-15 |
| CHIN | -0.7728 | 0.7324 | 1.5052 | 1.186e-15 | 5.987e-15 |
| Fragile histidine triad protein | -0.736 | 0.6592 | 1.3952 | 1.690e-15 | 8.430e-15 |
| Contactin-5 | -0.6678 | 0.8314 | 1.4991 | 1.841e-15 | 9.071e-15 |
| Calcineurin | -0.7615 | 0.8037 | 1.5652 | 2.598e-15 | 1.265e-14 |
| JAG1:ECD | -0.7437 | 0.7086 | 1.4522 | 3.062e-15 | 1.473e-14 |
| CAB39 | -0.7038 | 0.675 | 1.3788 | 3.581e-15 | 1.702e-14 |
| CG024 | -0.6768 | 0.7001 | 1.3769 | 4.383e-15 | 2.059e-14 |
| TCAL5 | -0.7157 | 0.7038 | 1.4196 | 4.603e-15 | 2.138e-14 |
| EFHD2 | -0.7345 | 0.7619 | 1.4964 | 9.130e-15 | 4.192e-14 |
| PTPA | -0.7146 | 0.6603 | 1.3749 | 1.541e-14 | 6.951e-14 |
| DAF | -0.6805 | 0.6831 | 1.3636 | 1.549e-14 | 6.951e-14 |
| NCAM-120 | -0.7031 | 0.7877 | 1.4908 | 2.047e-14 | 9.088e-14 |
| GLO2 | -0.6993 | 0.8026 | 1.502 | 2.329e-14 | 1.023e-13 |
| BOLA3 | -0.6889 | 0.7669 | 1.4558 | 3.141e-14 | 1.353e-13 |
| HSP 90b | -0.6969 | 0.6689 | 1.3658 | 3.147e-14 | 1.353e-13 |
| DLG4 | -0.6903 | 0.7129 | 1.4032 | 5.213e-14 | 2.217e-13 |
| PTPRR | -0.6768 | 0.8031 | 1.4798 | 5.885e-14 | 2.477e-13 |
| PPR1A | -0.7075 | 0.6864 | 1.3939 | 1.506e-13 | 6.272e-13 |
| COX5A | -0.6914 | 0.5328 | 1.2242 | 1.539e-13 | 6.344e-13 |
| AGAP3 | -0.6722 | 0.7159 | 1.3881 | 1.648e-13 | 6.724e-13 |
| DOC2B | -0.6969 | 0.6977 | 1.3946 | 2.053e-13 | 8.294e-13 |
| 14-3-3E | -0.6402 | 0.6931 | 1.3333 | 3.079e-13 | 1.232e-12 |
| LACB2 | -0.6547 | 0.6225 | 1.2772 | 3.275e-13 | 1.297e-12 |
| ARHL2 | -0.6585 | 0.6324 | 1.2909 | 5.569e-13 | 2.184e-12 |
| DNJB9 | -0.6966 | 0.7997 | 1.4964 | 8.028e-13 | 3.117e-12 |
| PCDH1 | -0.6753 | 0.8083 | 1.4836 | 8.101e-13 | 3.117e-12 |
| KIRR2 | -0.6958 | 0.6938 | 1.3896 | 8.439e-13 | 3.216e-12 |
| PKB gamma | -0.6848 | 0.678 | 1.3628 | 9.256e-13 | 3.495e-12 |
| PEX14:N-term | -0.6586 | 0.695 | 1.3536 | 1.215e-12 | 4.545e-12 |
| suppression of tumorigenicity 13 | -0.6695 | 0.6263 | 1.2958 | 1.542e-12 | 5.714e-12 |
| Nogo Receptor | -0.5527 | 0.715 | 1.2677 | 1.671e-12 | 6.137e-12 |
| PPM1B | -0.6589 | 0.6844 | 1.3433 | 1.773e-12 | 6.454e-12 |
| 7B2 | -0.6787 | 0.7646 | 1.4433 | 1.826e-12 | 6.588e-12 |
| CHKB | -0.6314 | 0.6093 | 1.2407 | 2.182e-12 | 7.802e-12 |
| PHIPL | -0.6633 | 0.6661 | 1.3293 | 2.837e-12 | 1.006e-11 |
| CRAC1 | -0.6828 | 0.6281 | 1.3109 | 3.661e-12 | 1.286e-11 |
| NUDT3 | -0.6412 | 0.6891 | 1.3303 | 3.965e-12 | 1.381e-11 |
| 14-3-3 eta | -0.6549 | 0.7089 | 1.3639 | 5.309e-12 | 1.833e-11 |
| PKHO2 | -0.5796 | 0.6974 | 1.277 | 5.834e-12 | 1.997e-11 |
| Nr-CAM | -0.6573 | 0.6196 | 1.277 | 6.564e-12 | 2.229e-11 |
| Glycyl t-RNA synthetase | -0.66 | 0.6678 | 1.3278 | 1.160e-11 | 3.906e-11 |
| CALB2 | -0.6848 | 0.6022 | 1.287 | 1.417e-11 | 4.731e-11 |
| GLNA | -0.6359 | 0.6646 | 1.3005 | 1.484e-11 | 4.916e-11 |
| HSP 90a | -0.6151 | 0.6475 | 1.2626 | 1.654e-11 | 5.420e-11 |
| TIPRL | -0.6515 | 0.5842 | 1.2357 | 1.663e-11 | 5.420e-11 |
| PGP9.5 | -0.6368 | 0.6769 | 1.3137 | 2.740e-11 | 8.855e-11 |
| ASB9 | -0.6544 | 0.67 | 1.3244 | 2.919e-11 | 9.361e-11 |
| Osteopontin | -0.6028 | 0.651 | 1.2538 | 3.652e-11 | 1.162e-10 |
| ISOC1 | -0.6156 | 0.6007 | 1.2163 | 4.094e-11 | 1.292e-10 |
| CLC2L | -0.6056 | 0.6631 | 1.2688 | 7.033e-11 | 2.203e-10 |
| VPS4A | -0.5814 | 0.734 | 1.3154 | 7.450e-11 | 2.315e-10 |
| COQ7 | 0.5784 | -0.6202 | -1.1986 | 7.885e-11 | 2.432e-10 |
| LRFN2 | -0.4641 | 0.69 | 1.1542 | 8.010e-11 | 2.451e-10 |
| NUDC:CS | -0.6559 | 0.5673 | 1.2232 | 8.441e-11 | 2.564e-10 |
| CB39L | -0.6443 | 0.5996 | 1.2439 | 1.394e-10 | 4.202e-10 |
| NEC1 | -0.5324 | 0.5923 | 1.1247 | 1.489e-10 | 4.457e-10 |
| EPN1 | -0.6438 | 0.5674 | 1.2112 | 1.569e-10 | 4.660e-10 |
| IF4B | -0.6279 | 0.5612 | 1.1891 | 1.717e-10 | 5.064e-10 |
| PCSK1 | -0.5984 | 0.7456 | 1.3439 | 2.079e-10 | 6.052e-10 |
| LIGO1 | -0.5045 | 0.7277 | 1.2322 | 2.082e-10 | 6.052e-10 |
| ARF1 | -0.5909 | 0.6325 | 1.2233 | 2.210e-10 | 6.378e-10 |
| F163B | -0.5986 | 0.6606 | 1.2591 | 2.400e-10 | 6.877e-10 |
| CCD50 | -0.6516 | 0.6072 | 1.2588 | 2.701e-10 | 7.683e-10 |
| Hemoglobin epsilon chain | -0.5917 | 0.7679 | 1.3596 | 3.855e-10 | 1.089e-09 |
| SPN90 | -0.5318 | 0.6636 | 1.1954 | 4.671e-10 | 1.310e-09 |
| VATF | -0.5735 | 0.5648 | 1.1383 | 4.883e-10 | 1.361e-09 |
| Cyclophilin A | -0.5904 | 0.5746 | 1.165 | 5.955e-10 | 1.648e-09 |
| DTD2 | -0.5868 | 0.6043 | 1.1911 | 6.703e-10 | 1.842e-09 |
| Radixin | -0.5802 | 0.5578 | 1.138 | 7.438e-10 | 2.030e-09 |
| CB032 | -0.5792 | 0.536 | 1.1153 | 8.709e-10 | 2.361e-09 |
| DCNL1 | -0.5701 | 0.6879 | 1.2579 | 9.884e-10 | 2.662e-09 |
| PHI | -0.5627 | 0.6372 | 1.1999 | 1.366e-09 | 3.644e-09 |
| MFGM | -0.5395 | 0.73 | 1.2695 | 1.371e-09 | 3.644e-09 |
| GGE2D | -0.5777 | 0.563 | 1.1407 | 1.923e-09 | 5.079e-09 |
| NEC2 | -0.5332 | 0.7988 | 1.3319 | 2.052e-09 | 5.383e-09 |
| ADH1B | -0.607 | 0.63 | 1.237 | 2.136e-09 | 5.567e-09 |
| STAR | -0.5989 | 0.621 | 1.2199 | 2.543e-09 | 6.586e-09 |
| RBM9 | -0.5702 | 0.5985 | 1.1687 | 3.146e-09 | 8.096e-09 |
| IGBP1 | -0.6022 | 0.5566 | 1.1588 | 3.996e-09 | 1.022e-08 |
| RWDD1 | -0.5618 | 0.5965 | 1.1583 | 4.367e-09 | 1.110e-08 |
| TXN4B | -0.5656 | 0.5961 | 1.1617 | 4.498e-09 | 1.136e-08 |
| LIGO3 | -0.5386 | 0.6572 | 1.1959 | 4.613e-09 | 1.157e-08 |
| WBP2 | -0.5556 | 0.6169 | 1.1725 | 4.724e-09 | 1.178e-08 |
| KRA24 | -0.5498 | 0.6945 | 1.2443 | 5.840e-09 | 1.447e-08 |
| CO9A3 | -0.5939 | 0.5197 | 1.1136 | 7.267e-09 | 1.790e-08 |
| UBC9 | -0.5507 | 0.5302 | 1.0808 | 7.452e-09 | 1.825e-08 |
| PDLI5 | -0.5811 | 0.4967 | 1.0778 | 9.594e-09 | 2.335e-08 |
| TNR6B | -0.5522 | 0.5423 | 1.0945 | 1.015e-08 | 2.456e-08 |
| NPTX1 | -0.4697 | 0.6916 | 1.1614 | 1.070e-08 | 2.573e-08 |
| RCD1 | -0.5804 | 0.5624 | 1.1428 | 1.205e-08 | 2.880e-08 |
| NT5D3 | -0.5644 | 0.5592 | 1.1237 | 1.473e-08 | 3.501e-08 |
| GAGE-2 | -0.5606 | 0.5642 | 1.1248 | 1.574e-08 | 3.718e-08 |
| IF4G2 | -0.5667 | 0.5321 | 1.0988 | 1.641e-08 | 3.854e-08 |
| SEM5B | -0.5308 | 0.5922 | 1.1229 | 2.132e-08 | 4.979e-08 |
| CNOT1 | -0.4762 | 0.5584 | 1.0346 | 2.155e-08 | 5.004e-08 |
| CCKN | -0.5391 | 0.6795 | 1.2186 | 2.864e-08 | 6.612e-08 |
| Alpha enolase | -0.5622 | 0.4952 | 1.0574 | 3.064e-08 | 7.033e-08 |
| UGPA1 | -0.5371 | 0.5818 | 1.1189 | 3.282e-08 | 7.492e-08 |
| CHSTA | -0.5145 | 0.651 | 1.1655 | 3.326e-08 | 7.548e-08 |
| SARP-2 | -0.4548 | 0.5512 | 1.0061 | 4.049e-08 | 9.139e-08 |
| PFD4 | -0.5076 | 0.4909 | 0.9985 | 4.550e-08 | 1.021e-07 |
| PGAM2 | -0.5517 | 0.5433 | 1.095 | 4.958e-08 | 1.107e-07 |
| MK08 | -0.5299 | 0.551 | 1.0809 | 6.199e-08 | 1.376e-07 |
| Corticotropin-lipotropin | -0.5356 | 0.4949 | 1.0305 | 6.337e-08 | 1.399e-07 |
| IRF1 | -0.3941 | 0.6876 | 1.0817 | 7.586e-08 | 1.666e-07 |
| ATX3 | -0.5766 | 0.5315 | 1.108 | 8.586e-08 | 1.875e-07 |
| RFNG | -0.5265 | 0.5497 | 1.0761 | 9.894e-08 | 2.149e-07 |
| CEI | -0.5253 | 0.6053 | 1.1307 | 1.074e-07 | 2.320e-07 |
| SCG1 | -0.5408 | 0.4489 | 0.9897 | 1.578e-07 | 3.391e-07 |
| KGUA | -0.5545 | 0.4459 | 1.0004 | 1.641e-07 | 3.509e-07 |
| SPC1L | -0.4693 | 0.6359 | 1.1053 | 1.826e-07 | 3.882e-07 |
| SMAP1 | -0.4767 | 0.6225 | 1.0992 | 2.262e-07 | 4.784e-07 |
| EMAP-2 | -0.4974 | 0.5132 | 1.0106 | 2.470e-07 | 5.198e-07 |
| Phospholipase C-delta-1 | -0.529 | 0.4565 | 0.9855 | 3.756e-07 | 7.861e-07 |
| HAX1 | -0.4747 | 0.558 | 1.0327 | 3.979e-07 | 8.287e-07 |
| FK506 binding protein 1A | -0.445 | 0.5696 | 1.0146 | 4.386e-07 | 9.088e-07 |
| IFNL4 | 0.5349 | -0.4579 | -0.9928 | 4.808e-07 | 9.910e-07 |
| SIA8A | -0.457 | 0.3635 | 0.8205 | 4.872e-07 | 9.991e-07 |
| CUTA | -0.4846 | 0.5201 | 1.0048 | 5.607e-07 | 1.144e-06 |
| EPHA4 | -0.4722 | 0.3846 | 0.8568 | 5.829e-07 | 1.183e-06 |
| PSB2 | -0.5001 | 0.5694 | 1.0694 | 6.333e-07 | 1.279e-06 |
| PDLI4 | -0.533 | 0.4159 | 0.9489 | 6.855e-07 | 1.378e-06 |
| SMOC2 | -0.515 | 0.3426 | 0.8576 | 7.029e-07 | 1.406e-06 |
| ISK2 | -0.5334 | 0.5106 | 1.044 | 7.646e-07 | 1.521e-06 |
| LIN7A | -0.4751 | 0.4777 | 0.9528 | 7.682e-07 | 1.521e-06 |
| TCAL3 | -0.4421 | 0.5872 | 1.0293 | 7.763e-07 | 1.530e-06 |
| PKB | -0.4659 | 0.4639 | 0.9298 | 9.307e-07 | 1.825e-06 |
| PIN1 | -0.4815 | 0.5221 | 1.0036 | 9.351e-07 | 1.825e-06 |
| UBXN4:CD 1 | -0.4808 | 0.4555 | 0.9363 | 9.639e-07 | 1.872e-06 |
| SYAP1 | -0.4826 | 0.4321 | 0.9147 | 9.754e-07 | 1.885e-06 |
| CAND1 | -0.5374 | 0.4582 | 0.9957 | 1.645e-06 | 3.165e-06 |
| CBLN2 | -0.4128 | 0.503 | 0.9158 | 2.206e-06 | 4.224e-06 |
| AHSA1 | -0.487 | 0.5124 | 0.9994 | 2.326e-06 | 4.433e-06 |
| SUMO4 | -0.4568 | 0.452 | 0.9088 | 2.385e-06 | 4.524e-06 |
| K0152 | -0.4698 | 0.5056 | 0.9754 | 2.412e-06 | 4.553e-06 |
| PFD2 | -0.4791 | 0.4573 | 0.9363 | 2.808e-06 | 5.277e-06 |
| PAI-2 | -0.4562 | 0.4617 | 0.918 | 3.431e-06 | 6.417e-06 |
| FKBP52 protein | -0.4823 | 0.4769 | 0.9592 | 3.538e-06 | 6.587e-06 |
| LRRF2 | -0.4622 | 0.457 | 0.9192 | 3.728e-06 | 6.878e-06 |
| ULBP-2 | -0.3878 | 0.522 | 0.9098 | 3.728e-06 | 6.878e-06 |
| SYT12 | -0.4807 | 0.4165 | 0.8972 | 4.392e-06 | 8.065e-06 |
| MGT4B | -0.5077 | 0.4277 | 0.9353 | 5.879e-06 | 1.075e-05 |
| Integrin a1b1 | -0.4275 | 0.5032 | 0.9308 | 6.151e-06 | 1.119e-05 |
| PEA15 | -0.4403 | 0.5327 | 0.973 | 6.329e-06 | 1.147e-05 |
| MCEE | -0.4276 | 0.4577 | 0.8854 | 6.432e-06 | 1.160e-05 |
| CCYL1 | -0.456 | 0.3958 | 0.8519 | 7.396e-06 | 1.328e-05 |
| PROSC | -0.4296 | 0.5216 | 0.9512 | 7.464e-06 | 1.334e-05 |
| UBE2H | -0.3877 | 0.5244 | 0.9121 | 7.553e-06 | 1.344e-05 |
| MEK1 | -0.3923 | 0.5021 | 0.8944 | 7.628e-06 | 1.352e-05 |
| SYWC | -0.4705 | 0.4458 | 0.9163 | 8.767e-06 | 1.547e-05 |
| CAMLG | -0.4586 | 0.5052 | 0.9638 | 9.018e-06 | 1.584e-05 |
| Proteasome beta chain | -0.374 | 0.4993 | 0.8733 | 1.375e-05 | 2.405e-05 |
| LRRN1:CD | -0.4297 | 0.4941 | 0.9238 | 1.474e-05 | 2.566e-05 |
| HOME1 | -0.4458 | 0.4009 | 0.8467 | 1.496e-05 | 2.594e-05 |
| MPC1 | -0.3944 | 0.4706 | 0.865 | 1.736e-05 | 2.998e-05 |
| PSA7 | -0.4039 | 0.3848 | 0.7887 | 2.641e-05 | 4.541e-05 |
| RIC8A | -0.4291 | 0.4279 | 0.857 | 3.318e-05 | 5.681e-05 |
| CMTD1 | -0.4703 | 0.3703 | 0.8406 | 3.347e-05 | 5.705e-05 |
| MK10 | -0.4607 | 0.3925 | 0.8531 | 3.595e-05 | 6.102e-05 |
| Histidyl-tRNA synthetase | -0.4235 | 0.4079 | 0.8314 | 3.885e-05 | 6.567e-05 |
| GGA3 | -0.3702 | 0.4668 | 0.837 | 4.380e-05 | 7.373e-05 |
| FGF-19 | -0.4462 | 0.3056 | 0.7518 | 5.515e-05 | 9.244e-05 |
| PPME1 | -0.4171 | 0.4612 | 0.8784 | 5.625e-05 | 9.391e-05 |
| BN3D2 | -0.3973 | 0.4524 | 0.8496 | 6.088e-05 | 1.012e-04 |
| RND1 | -0.3596 | 0.4615 | 0.8211 | 6.860e-05 | 1.136e-04 |
| TTC9A | -0.4182 | 0.4149 | 0.8331 | 7.734e-05 | 1.275e-04 |
| OTUL | -0.3489 | 0.4438 | 0.7927 | 8.466e-05 | 1.390e-04 |
| Cystatin B | -0.4189 | 0.2923 | 0.7112 | 8.986e-05 | 1.470e-04 |
| NCK2 | -0.3742 | 0.4199 | 0.7942 | 1.123e-04 | 1.830e-04 |
| SPC25 | -0.3682 | 0.4611 | 0.8292 | 1.187e-04 | 1.925e-04 |
| Myokinase, human | -0.4614 | 0.2992 | 0.7606 | 1.206e-04 | 1.948e-04 |
| RN215:ECD | -0.3511 | 0.3827 | 0.7338 | 1.290e-04 | 2.076e-04 |
| 2A5A | -0.3682 | 0.4061 | 0.7743 | 1.305e-04 | 2.092e-04 |
| F209B | -0.3651 | 0.3707 | 0.7359 | 1.313e-04 | 2.096e-04 |
| PHP14 | -0.4115 | 0.4283 | 0.8398 | 1.384e-04 | 2.201e-04 |
| TXNDC4 | -0.4588 | 0.255 | 0.7138 | 1.566e-04 | 2.482e-04 |
| Phosphoglycerate mutase 1 | -0.3225 | 0.4772 | 0.7997 | 1.715e-04 | 2.706e-04 |
| TATD3 | -0.2525 | 0.4828 | 0.7353 | 1.730e-04 | 2.720e-04 |
| CHL1 | -0.3621 | 0.3728 | 0.7349 | 1.823e-04 | 2.855e-04 |
| p15-INK4b | -0.4044 | 0.3517 | 0.756 | 2.005e-04 | 3.128e-04 |
| HSP70 protein 8 | -0.3916 | 0.3762 | 0.7678 | 2.231e-04 | 3.467e-04 |
| FA69C | -0.3763 | 0.3073 | 0.6836 | 2.953e-04 | 4.571e-04 |
| Siglec-7 | -0.4332 | 0.2662 | 0.6993 | 3.262e-04 | 5.029e-04 |
| PSA2 | -0.3344 | 0.3705 | 0.705 | 3.300e-04 | 5.069e-04 |
| ARL1 | -0.3214 | 0.3521 | 0.6734 | 3.650e-04 | 5.585e-04 |
| Apo E2 | 0.4045 | -0.2775 | -0.682 | 3.893e-04 | 5.934e-04 |
| Nucleoside diphosphate kinase A | -0.3337 | 0.3869 | 0.7206 | 4.305e-04 | 6.538e-04 |
| G3BP2 | -0.3492 | 0.3241 | 0.6732 | 4.529e-04 | 6.852e-04 |
| AGR2 | -0.3534 | 0.3421 | 0.6955 | 6.464e-04 | 9.744e-04 |
| TMOD2 | -0.3415 | 0.3468 | 0.6883 | 6.499e-04 | 9.760e-04 |
| TRIML | 0.3329 | -0.3561 | -0.689 | 7.566e-04 | 1.132e-03 |
| SH3G2 | -0.4132 | 0.437 | 0.8502 | 8.565e-04 | 1.277e-03 |
| HBAT | -0.3644 | 0.3071 | 0.6715 | 9.295e-04 | 1.381e-03 |
| NFL | -0.3634 | 0.2439 | 0.6073 | 1.136e-03 | 1.680e-03 |
| PAFAH beta subunit | -0.4146 | 0.1362 | 0.5508 | 1.459e-03 | 2.151e-03 |
| STRATIFIN | -0.3516 | 0.3022 | 0.6538 | 1.687e-03 | 2.479e-03 |
| HXK1 | -0.2711 | 0.3742 | 0.6453 | 1.772e-03 | 2.594e-03 |
| FOPNL | -0.2748 | 0.3582 | 0.633 | 2.173e-03 | 3.169e-03 |
| NFH | -0.3537 | 0.1962 | 0.5499 | 2.236e-03 | 3.249e-03 |
| RhoGDI | -0.2933 | 0.4478 | 0.7411 | 2.519e-03 | 3.647e-03 |
| 2A5D | -0.2346 | 0.4298 | 0.6644 | 2.840e-03 | 4.098e-03 |
| NUDT7 | -0.3277 | 0.2713 | 0.5991 | 3.185e-03 | 4.580e-03 |
| DPYL5 | -0.2611 | 0.3836 | 0.6447 | 3.246e-03 | 4.650e-03 |
| CK5P3 | -0.2738 | 0.2517 | 0.5255 | 3.447e-03 | 4.921e-03 |
| P5I11 | -0.3563 | 0.2442 | 0.6005 | 3.579e-03 | 5.091e-03 |
| NUPL | -0.3088 | 0.2765 | 0.5853 | 3.980e-03 | 5.634e-03 |
| HID1 | -0.3109 | 0.2551 | 0.5659 | 3.989e-03 | 5.634e-03 |
| MMP-10 | -0.3378 | 0.2268 | 0.5646 | 4.542e-03 | 6.393e-03 |
| PH | 0.2349 | -0.3546 | -0.5895 | 4.713e-03 | 6.611e-03 |
| NIP7 | 0.1879 | -0.3404 | -0.5283 | 5.471e-03 | 7.648e-03 |
| TPPC3 | -0.3241 | 0.1829 | 0.5069 | 6.193e-03 | 8.609e-03 |
| MIF | -0.2849 | 0.3041 | 0.589 | 6.201e-03 | 8.609e-03 |
| ZN483 | -0.2955 | 0.2294 | 0.5249 | 6.390e-03 | 8.840e-03 |
| MK01 | -0.2014 | 0.3587 | 0.5601 | 6.556e-03 | 9.040e-03 |
| RAB13 | -0.3478 | 0.2303 | 0.578 | 6.599e-03 | 9.069e-03 |
| GNAI3 | -0.2314 | 0.326 | 0.5574 | 6.657e-03 | 9.117e-03 |
| NECA3 | -0.3108 | 0.259 | 0.5697 | 7.048e-03 | 9.619e-03 |
| IRF6 | -0.3134 | 0.1933 | 0.5067 | 7.158e-03 | 9.736e-03 |
| KCTD2 | -0.2487 | 0.3031 | 0.5518 | 8.291e-03 | 1.124e-02 |
| TBCE | -0.2693 | 0.2681 | 0.5374 | 8.340e-03 | 1.127e-02 |
| DCNL5 | -0.3401 | 0.1324 | 0.4725 | 9.027e-03 | 1.216e-02 |
| EIF1A | -0.2706 | 0.2423 | 0.5129 | 1.020e-02 | 1.369e-02 |
| FLII | -0.2439 | 0.272 | 0.5158 | 1.038e-02 | 1.388e-02 |
| PSB6 | -0.2777 | 0.2474 | 0.5251 | 1.135e-02 | 1.509e-02 |
| CLUL1 | -0.2447 | 0.2756 | 0.5203 | 1.139e-02 | 1.509e-02 |
| SELS | -0.2933 | 0.2277 | 0.521 | 1.139e-02 | 1.509e-02 |
| ODPX | -0.2576 | 0.2602 | 0.5179 | 1.150e-02 | 1.519e-02 |
| C56D1 | 0.3269 | -0.1897 | -0.5166 | 1.197e-02 | 1.575e-02 |
| sTREM-1 | -0.22 | 0.3162 | 0.5362 | 1.228e-02 | 1.611e-02 |
| ILT-4 | -0.2616 | 0.221 | 0.4826 | 1.314e-02 | 1.719e-02 |
| KAT | -0.2431 | 0.2623 | 0.5054 | 1.428e-02 | 1.862e-02 |
| OPG | -0.2876 | 0.1352 | 0.4228 | 1.526e-02 | 1.982e-02 |
| TECK | -0.2252 | 0.2567 | 0.4819 | 1.681e-02 | 2.176e-02 |
| EFS | 0.185 | -0.2531 | -0.4382 | 2.011e-02 | 2.596e-02 |
| LX15B | 0.2389 | -0.2229 | -0.4618 | 2.289e-02 | 2.944e-02 |
| NCOA7 | -0.1812 | 0.2818 | 0.463 | 2.385e-02 | 3.059e-02 |
| CCD24 | -0.2164 | 0.2823 | 0.4988 | 2.434e-02 | 3.112e-02 |
| FOXO1A | -0.2596 | 0.1667 | 0.4263 | 2.459e-02 | 3.134e-02 |
| MOBL3 | -0.2486 | 0.1897 | 0.4382 | 2.619e-02 | 3.327e-02 |
| THY1 | -0.1476 | 0.3019 | 0.4495 | 2.827e-02 | 3.581e-02 |
| PSMA | -0.163 | 0.3055 | 0.4685 | 2.855e-02 | 3.605e-02 |
| CAMK2B | -0.1423 | 0.306 | 0.4483 | 3.033e-02 | 3.817e-02 |
| QORL1 | -0.2631 | 0.174 | 0.4371 | 3.349e-02 | 4.202e-02 |
| Albumin | 0.1404 | -0.2788 | -0.4192 | 3.778e-02 | 4.725e-02 |
| Keratin 19 | -0.1998 | 0.2397 | 0.4395 | 4.033e-02 | 5.029e-02 |
| Glypican 3 | 0.1849 | -0.217 | -0.4019 | 4.587e-02 | 5.701e-02 |
| PICK1 | -0.2129 | 0.1756 | 0.3885 | 4.666e-02 | 5.782e-02 |
| Secretagogin | -0.2186 | 0.25 | 0.4686 | 4.969e-02 | 6.139e-02 |
| ARL2 | -0.2221 | 0.1798 | 0.4019 | 5.073e-02 | 6.248e-02 |
| MTMR7 | -0.227 | 0.1397 | 0.3668 | 5.146e-02 | 6.319e-02 |
| LMO4 | -0.2066 | 0.1913 | 0.3979 | 5.536e-02 | 6.778e-02 |
| RUXF | -0.2455 | 0.118 | 0.3636 | 5.945e-02 | 7.257e-02 |
| BORC5 | -0.1663 | 0.2202 | 0.3865 | 6.281e-02 | 7.643e-02 |
| CRYL1 | -0.184 | 0.1889 | 0.3729 | 6.900e-02 | 8.372e-02 |
| K0494:C-term | -0.1545 | 0.2437 | 0.3982 | 6.926e-02 | 8.377e-02 |
| AP1B1 | -0.1511 | 0.2235 | 0.3746 | 6.955e-02 | 8.387e-02 |
| TATD1 | -0.2087 | 0.1447 | 0.3535 | 7.557e-02 | 9.086e-02 |
| PNMA2 | 0.2152 | -0.1498 | -0.365 | 7.643e-02 | 9.162e-02 |
| R3GEF | -0.131 | 0.236 | 0.367 | 7.862e-02 | 9.397e-02 |
| S100A16 | -0.2278 | 0.1025 | 0.3304 | 8.966e-02 | 1.069e-01 |
| Galectin-1 | -0.1973 | 0.1455 | 0.3428 | 9.089e-02 | 1.080e-01 |
| Chitotriosidase-1 | -0.1901 | 0.1446 | 0.3347 | 9.362e-02 | 1.109e-01 |
| ATG7 | 0.1609 | -0.1855 | -0.3465 | 9.461e-02 | 1.118e-01 |
| SPD2B | -0.1298 | 0.2222 | 0.352 | 9.781e-02 | 1.152e-01 |
| SNPH | 0.2302 | -0.098 | -0.3282 | 1.008e-01 | 1.184e-01 |
| VCIP1 | 0.0943 | -0.2344 | -0.3287 | 1.108e-01 | 1.298e-01 |
| SHAN1 | -0.1422 | 0.1773 | 0.3194 | 1.120e-01 | 1.308e-01 |
| IMA6 | -0.2097 | 0.0742 | 0.2839 | 1.275e-01 | 1.484e-01 |
| TMA16 | -0.1504 | 0.1677 | 0.3181 | 1.322e-01 | 1.534e-01 |
| TBCA | -0.1458 | 0.1831 | 0.3289 | 1.381e-01 | 1.599e-01 |
| PP2AAA | -0.1874 | 0.11 | 0.2973 | 1.385e-01 | 1.599e-01 |
| GLRX3 | -0.2309 | 0.0623 | 0.2932 | 1.673e-01 | 1.925e-01 |
| CF6 | -0.1783 | 0.1058 | 0.2841 | 1.718e-01 | 1.972e-01 |
| MAGD1 | -0.1445 | 0.1212 | 0.2657 | 1.804e-01 | 2.064e-01 |
| RNF43 | 0.1675 | -0.0802 | -0.2477 | 1.869e-01 | 2.134e-01 |
| PNOC | -0.1421 | 0.1252 | 0.2674 | 1.929e-01 | 2.195e-01 |
| ATE1 | -0.1697 | 0.0955 | 0.2652 | 1.948e-01 | 2.210e-01 |
| SMAD5 | -0.092 | 0.1617 | 0.2537 | 2.019e-01 | 2.285e-01 |
| Cofilin-1 | -0.0603 | 0.2156 | 0.2759 | 2.046e-01 | 2.309e-01 |
| Apo B | 0.1429 | -0.1146 | -0.2575 | 2.129e-01 | 2.396e-01 |
| CINP | -0.0601 | 0.1907 | 0.2509 | 2.149e-01 | 2.412e-01 |
| HCC-1 | -0.1326 | 0.1025 | 0.2351 | 2.199e-01 | 2.461e-01 |
| ZRAN1 | -0.1265 | 0.1155 | 0.242 | 2.245e-01 | 2.505e-01 |
| Angiotensinogen | 0.1382 | -0.1014 | -0.2396 | 2.311e-01 | 2.572e-01 |
| MAAI | 0.1002 | -0.1292 | -0.2294 | 2.385e-01 | 2.648e-01 |
| RDH16 | -0.1451 | 0.0703 | 0.2154 | 2.544e-01 | 2.816e-01 |
| CONA1 | 0.0922 | -0.1291 | -0.2213 | 2.655e-01 | 2.930e-01 |
| VPS29 | 0.1075 | -0.1029 | -0.2104 | 2.914e-01 | 3.208e-01 |
| CHFR | 0.1381 | -0.0581 | -0.1962 | 2.929e-01 | 3.216e-01 |
| ISK13 | -0.115 | 0.0961 | 0.2111 | 2.973e-01 | 3.255e-01 |
| GON1 | 0.0285 | -0.188 | -0.2165 | 3.122e-01 | 3.409e-01 |
| S100A13 | -0.1066 | 0.1145 | 0.2212 | 3.134e-01 | 3.413e-01 |
| alpha-1-antichymotrypsin complex | -0.1233 | 0.0583 | 0.1816 | 3.358e-01 | 3.647e-01 |
| HS105 | -0.0438 | 0.1469 | 0.1907 | 3.568e-01 | 3.864e-01 |
| WN10B | -0.1327 | 0.0457 | 0.1783 | 3.839e-01 | 4.147e-01 |
| NFIA | 0.1076 | -0.0436 | -0.1512 | 4.300e-01 | 4.632e-01 |
| PPIL1 | -0.1364 | 0.0237 | 0.1602 | 4.325e-01 | 4.647e-01 |
| MAZ | -0.0947 | 0.0603 | 0.155 | 4.423e-01 | 4.740e-01 |
| CAMK2D | 0.1568 | -0.0015 | -0.1583 | 4.438e-01 | 4.743e-01 |
| FSD1 | -0.0559 | 0.0939 | 0.1497 | 4.691e-01 | 5.000e-01 |
| SIRT5 | -0.0392 | 0.1127 | 0.1519 | 4.851e-01 | 5.157e-01 |
| Ataxin-2-binding protein 1 | -0.1586 | -0.019 | 0.1396 | 5.074e-01 | 5.380e-01 |
| CHCH7 | 0.0021 | 0.116 | 0.1139 | 5.448e-01 | 5.762e-01 |
| NEP1 | -0.0539 | 0.0667 | 0.1206 | 5.676e-01 | 5.987e-01 |
| NUD11 | -0.0867 | 0.0219 | 0.1087 | 5.720e-01 | 6.018e-01 |
| CN37 | 0.1056 | -0.0042 | -0.1098 | 6.042e-01 | 6.340e-01 |
| IFIT2 | -0.0054 | -0.1039 | -0.0986 | 6.226e-01 | 6.517e-01 |
| PTPM1 | -0.1174 | -0.0193 | 0.0981 | 6.331e-01 | 6.609e-01 |
| IMA5 | 0.0308 | -0.0684 | -0.0992 | 6.412e-01 | 6.677e-01 |
| MAGE-3 | -0.0891 | -0.0123 | 0.0768 | 6.796e-01 | 7.058e-01 |
| RBBP4 | -0.0202 | 0.0541 | 0.0743 | 7.066e-01 | 7.319e-01 |
| UCHL5 | -0.0541 | 0.0234 | 0.0775 | 7.089e-01 | 7.325e-01 |
| Trypsin | -0.0212 | 0.0484 | 0.0697 | 7.320e-01 | 7.545e-01 |
| AK1C4 | -0.0294 | 0.043 | 0.0724 | 7.369e-01 | 7.575e-01 |
| ARL8A | 0.0215 | -0.0363 | -0.0578 | 7.624e-01 | 7.818e-01 |
| PTPRN | -0.0506 | -0.0125 | 0.0382 | 8.416e-01 | 8.608e-01 |
| HYKK | -0.0274 | -0.0657 | -0.0384 | 8.528e-01 | 8.700e-01 |
| DPEP1 | 0.0041 | -0.0345 | -0.0386 | 8.573e-01 | 8.724e-01 |
| SENP8 | -0.0001 | 0.0344 | 0.0345 | 8.720e-01 | 8.851e-01 |
| DNJA4 | -0.025 | -0.0556 | -0.0305 | 8.743e-01 | 8.852e-01 |
| SULT 1E | 0.0308 | -0.0032 | -0.034 | 8.790e-01 | 8.878e-01 |
| CABP7 | -0.0417 | -0.066 | -0.0243 | 9.071e-01 | 9.139e-01 |
| SDF2 | 0.025 | 0.0456 | 0.0205 | 9.210e-01 | 9.255e-01 |
| MVK | -0.0478 | -0.0627 | -0.0149 | 9.443e-01 | 9.466e-01 |
| MIRO1 | 0.0345 | 0.0265 | -0.008 | 9.699e-01 | 9.699e-01 |

*Abbreviations: FDR, false discovery rate (Benjamini–Hochberg); Sub., Subtype.*

## Notes

### Competing Interest Statement

The authors have declared no competing interest.

## REFERENCES

1. Jack CR Jr, Bennett DA, Blennow K, et al. NIA-AA Research Framework: Toward a biological definition of Alzheimer’s disease. Alzheimer’s & Dementia. 2018;14(4):535–562. 10.1016/j.jalz.2018.02.018

2. De Strooper B, Karran E. The cellular phase of Alzheimer’s disease. Cell. 2016;164(4):603–615. 10.1016/j.cell.2015.12.056

3. Karran E, De Strooper B. The amyloid cascade hypothesis: are we poised for success or failure? Journal of Neurochemistry. 2016;139(Suppl 2):237–252. 10.1111/jnc.13632

4. Selkoe DJ. Alzheimer’s disease is a synaptic failure. Science. 2002;298(5594):789–791. 10.1126/science.1074069

5. Duara R, Barker W. Heterogeneity in Alzheimer’s disease diagnosis and progression rates:Implications for Therapeutic Trials. Neurotherapeutics. 2022;19(1):8–25. 10.1007/s13311-022-01185-z

6. Ferreira D, et al. Biological subtypes of Alzheimer disease: A systematic review and meta-analysis. Neurology. 2020;94(10):436–448. 10.1212/WNL.0000000000009058

7. Yi, Luopeiwen, et al. Topological and geometric signatures of brain network dynamics in Alzheimer’s disease. Alzheimer’s & Dementia 21.8 (2025): e70545.

8. Wang M, et al. The Mount Sinai cohort of large-scale genomic, transcriptomic and proteomic data in Alzheimer’s disease. Scientific Data. 2018;5:180185. 10.1038/sdata.2018.185

9. Zhang B, et al. Integrated systems approach identifies genetic nodes and networks in late-onset Alzheimer’s disease. Cell. 2013;153(3):707–720. 10.1016/j.cell.2013.03.030

10. Sperling RA, et al. Toward defining the preclinical stages of Alzheimer’s disease. Alzheimer’s & Dementia. 2011;7(3):280–292. 10.1016/j.jalz.2011.03.003

11. Hou X-H, Zhang W, Kang K, et al. Proteomics reveals three molecular subtypes of Alzheimer’s disease with distinct progression patterns. Alzheimer’s & Dementia. 2026;22:e71106. 10.1002/alz.71106

12. The Gene Ontology Consortium. The Gene Ontology resource: enriching a GOld mine. Nucleic Acids Research. 2021;49(D1):D325–D334. 10.1093/nar/gkaa1113

13. Lambert JC, et al. Meta-analysis of 74,046 individuals identifies 11 new susceptibility loci for Alzheimer’s disease. Nature Genetics. 2013;45(12):1452–1458. 10.1038/ng.2802

14. Braak H, Braak E. Neuropathological stageing of Alzheimer-related changes. Acta Neuropathologica. 1991;82(4):239–259. 10.1007/BF00308809

15. Villemagne VL, et al. Amyloid beta deposition, neurodegeneration, and cognitive decline. Lancet Neurology. 2013;12(4):357–367. 10.1016/S1474-4422(13)70044-9

16. Avelar-Pereira B, Belloy ME, O’Hara R, Hosseini SMH. Decoding the heterogeneity of Alzheimer’s disease diagnosis and progression using multilayer networks. Molecular Psychiatry. 2023;28(6):2423–2432. 10.1038/s41380-022-01886-z

17. Vapnik VN. Statistical Learning Theory. New York, NY: Wiley; 1998. ISBN: 978-0-471-03003-4.

18. Tijms BM, et al. Cerebrospinal fluid proteomics in patients with Alzheimer’s disease reveals five molecular subtypes with distinct genetic risk profiles. Nature Aging. 2024;4:33–47. 10.1038/s43587-023-00550-7

19. Coifman RR, Lafon S. Diffusion maps. Applied and Computational Harmonic Analysis. 2006;21(1):5–30. 10.1016/j.acha.2006.04.006

20. Edelsbrunner H, Harer J. Computational Topology: An Introduction. American Mathematical Society; 2010. ISBN: 978-0-8218-4925-5

21. Edelsbrunner H, Harer J. Persistent homology—a survey. Contemporary Mathematics. 2008;453:257–282. 10.1090/conm/453

22. Singh G, et al. Topological methods for analysis of high dimensional data sets and 3D object recognition. Eurographics Symposium on Point-Based Graphics. 2007:91–100. 10.2312/SPBG/SPBG07/091-100

23. Liu FT, Ting KM, Zhou ZH. Isolation Forest. IEEE International Conference on Data Mining. 2008:413–422. 10.1109/ICDM.2008.17

24. Filzmoser P, Maronna R, Werner M. Outlier identification in high dimensions. Computational Statistics & Data Analysis. 2008;52(3):1694– 1711. 10.1016/j.csda.2007.05.018

25. Chazal F, Michel B. An introduction to topological data analysis: fundamental and practical aspects for data scientists. arXiv. 2017. https://arxiv.org/abs/1710.04019

26. Songdechakraiwut, Tananun, Chung Moo K. Dynamic topological data analysis for functional brain signals. International Symposium on Biomedical Imaging (ISBI). IEEE, 2020.

27. Lee H, Kang H, Chung MK, Kim B-N, Lee DS. Persistent brain network homology from the perspective of dendrogram. IEEE Transactions on Medical Imaging. 2012;31(12):2267–2277. 10.1109/TMI.2012.2219590

28. Kruskal JB. On the shortest spanning subtree of a graph and the traveling salesman problem. Proceedings of the American Mathematical Society. 1956;7(1):48–50. 10.1090/S0002-9939-1956-0078686-7

29. Skraba P, Turner K. Wasserstein stability for persistence diagrams. *arXiv preprint arXiv:2006.16824*. 2020. https://arxiv.org/abs/2006.16824

30. Raudvere U, et al. g:Profiler: a web server for functional enrichment analysis and conversions of gene lists. Nucleic Acids Research. 2019;47(W1):W191–W198. 10.1093/nar/gkz369

31. Jassal B, et al. The Reactome pathway knowledgebase. Nucleic Acids Research. 2020;48(D1):D498–D503. 10.1093/nar/gkz1031

32. Giusti C, et al. Clique topology reveals intrinsic geometric structure in neural correlations. PNAS. 2015;112(44):13455–13460. 10.1073/pnas.1506407112

33. Carlsson G. Topology and data. Bulletin of the American Mathematical Society. 2009;46(2):255–308. 10.1090/S0273-0979-09-01249-X

34. Munkres JR. Elements of Algebraic Topology. Boca Raton, FL: CRC Press; 2018. 10.1201/9780429493911

35. Songdechakraiwut T, Chung MK. Topological learning for brain networks. The Annals of Applied Statistics 2023;17(1):403–433. 10.1214/22-AOAS1633

36. Nicolau M, Levine AJ, Carlsson G. Topology based data analysis identifies a subgroup of breast cancers with a unique mutational profile and excellent survival. Proceedings of the National Academy of Sciences. 2011;108(17):7265–7270. 10.1073/pnas.1102826108

37. Sporns O. Network neuroscience. Nature Neuroscience. 2013. 2017;20:353–364. 10.1038/nn.4502

